# Membrane affinity difference between MinD monomer and dimer is not crucial for MinD gradient formation in Bacillus subtilis

**DOI:** 10.1101/2024.06.11.598461

**Authors:** Laura C. Bohorquez, Henrik Strahl, Davide Marenduzzo, Martin J. Thiele, Frank Bürmann, Leendert W. Hamoen

**Affiliations:** The Bacterial Cell Biology, Swammerdam Institute for Life Sciences, University of Amsterdam, Amsterdam, The Netherlands; The Centre for Bacterial Cell Biology, Biosciences Institute, Newcastle University, Newcastle upon Tyne, United Kingdom; SUPA, School of Physics and Astronomy, The University of Edinburgh, Edinburgh, Scotland, United Kingdom

## Abstract

Proteins can diffuse micrometers in seconds, yet bacterial cells are able to maintain stable protein gradients. The best studied bacterial protein gradient is the Min system of Escherichia coli. In rod-shaped bacteria the MinCD proteins prevent formation of minicells by inhibiting FtsZ polymerization close to the cell poles. In E. coli these proteins oscillate between cell poles within a minute, resulting in an increased MinCD concentration at the poles. This oscillation is caused by the interaction between MinD and the protein MinE, which form an ATP-driven reaction-diffusion system, whereby the ATPase MinD cycles between a monomeric cytosolic and a dimeric membrane attached states. Bacillus subtilis also has MinCD, but lacks MinE. In this case MinCD form a static gradient that requires the transmembrane protein MinJ, located at cell poles and cell division sites. A recent reaction-diffusion model was successful in recreating the MinD gradient in B. subtilis, assuming that MinD cycles between cytosol and membrane, like in E. coli. Here we show that the monomeric and dimeric states of B. subtilis MinD have comparable membrane affinities, that MinD interacts with MinJ as a dimer, and that MinJ is not required for membrane localization of MinD. Based on these new findings we tested different models, using kinetic Monte Carlo simulations, and found that a difference in diffusion rate between the monomer and dimer, rather than a difference in membrane affinity, is important for B. subtilis MinCD gradient formation.

## INTRODUCTION

Bacterial cells are capable of positioning proteins at midcell and cell poles, and can form protein gradients without the support of specific membrane compartments or guidance by cytoskeleton elements, as found in eukaryotic cells. A well-known example is the formation of a MinCD protein gradient along the cell axis of rod-shaped E. coli and B. subtilis cells. These proteins ensure that cell division occurs at midcell and not close to cell poles (1). The formation of these protein gradients is remarkable considering that the diffusion of proteins in cells is in the order of μm square per second, whereas these bacteria are only a few µm in length (2).

The Min system of E. coli is the best studied gradient system, and comprises the membrane associated MinD and MinE proteins that form a reaction-diffusion couple (3–5), resulting in a MinCD gradient that oscillates between cell poles in a matter of seconds (6, 7). This system has been reconstructed on artificial membranes resulting in dynamic wave patterns (8), and extensively simulated to understand the reaction parameters (e.g. (3–5, 9–12)). B. subtilis also forms a MinD gradient that decreases in concentration from cell poles and nascent cell division sites towards the middle of the cell. However, unlike E. coli, this gradient does not oscillate (13–15). In this study, we investigated how the MinCD gradient is formed in B. subtilis.

The core of the Min system consists of the protein couple MinC and MinD that prevent aberrant polymerization of the key cell division protein FtsZ close to newly formed septa or cell poles (15–17). MinC inhibits polymerization of FtsZ by direct protein-protein interactions and needs to bind to the Walker A-type ATPase MinD for its recruitment to the polar regions of the cell (18–21). Binding of ATP leads to MinD dimerization (22, 23), and subsequent association with the cell membrane, whereby its C-terminal amphipathic helix functions as membrane anchor (23, 24). MinC is recruited to the cell membrane by association with MinD dimers (23, 25, 26). The E. coli Min system comprises a third protein, the peripheral membrane protein MinE, that interacts with MinD, thereby displacing MinC and stimulating the ATPase activity of MinD, which ultimately triggers the release of MinD from the membrane (25, 27–30). The interaction with MinD causes a conformational change in MinE that stimulates its membrane affinity (27). These reciprocal interactions of MinD and MinE represent a natural reaction-diffusion system, resulting in the oscillation of MinCD proteins between cell poles with a periodicity of approximately 50 seconds (6–8). Through this oscillation, the average concentration of MinC is higher close to cell poles and lower at midcell, thereby inhibiting polymerization of FtsZ close to cell poles and favoring Z-ring formation at midcell (31).

The MinCD proteins of E. coli and B. subtilis are highly conserved. However, B. subtilis does not encode a MinE homologue and instead requires the proteins DivIVA and MinJ for the proper polar and septal localization of MinCD (32, 33). DivIVA is a general scaffold protein conserved in Gram-positive bacteria that associates with strong negatively curved membrane areas at the base of division septa, resulting in polarly located DivIVA clusters after division is completed (34, 35). The integral membrane protein MinJ interacts with DivIVA and shows the same localization pattern. MinJ also interacts with MinD, and inactivation of MinJ eliminates a MinCD gradient, resulting in filamentous cells and minicell formation due to uncontrolled activity of MinC (32, 33). Since DivIVA is recruited to midcell in the early stages of septum synthesis when a strong concave membrane area is formed (34), the Min proteins are already present when septum synthesis is ongoing, and their primary activity it to inhibit Z-ring formation next to nascent division sites (15).

The ATP-driven cycle between membrane association and dissociation of MinD is critical to the formation of the oscillating MinD gradient in E. coli. Mathematical modelling has shown that such an ATP-driven reversible membrane attachment cycle can also explain the formation of a MinD gradient in B. subtilis (36, 37). However, the C-terminal amphipathic alpha helix of B. subtilis MinD has a much stronger affinity for the cell membrane compared to its E. coli counterpart, and when fused to GFP can recruit this protein to the cell membrane, which the E. coli MinD C-terminal membrane anchor is not capable of (24). In fact, a MinD point mutant that prevents binding of ATP and maintains the protein in its monomeric state is still recruited to the B. subtilis cell membrane (38). It is therefore unclear whether dimerization-dependent changes in the membrane affinity of MinD are crucial for the formation of a B. subtilis MinCD gradient.

In this study, we analyzed and manipulated the membrane affinity of B. subtilis MinD, and combined in vivo experiments with in silico Monte Carlo simulations to assess the role of membrane affinity in the formation of a MinCD gradient. The main difference between our kinetic Monte Carlo model and previous mathematic models based on reaction-diffusion equation is that our treatment is particle-based, rather than continuum field-based, and naturally includes the effect of stochastic fluctuations, i.e. noise of the system. The experiments demonstrated that, unlike in E. coli, a difference in membrane affinity between MinD monomer and dimer is not critical for the creation of a MinCD gradient. In addition, we examined how differences in MinD membrane affinity affect the recruitment of MinC to the membrane, a process that is critical for the FtsZ-regulating activity of the Min-system.

## RESULTS

### Effect of dimerization on the localization of B. subtilis MinD

We began by examining how much the membrane affinity of monomeric B. subtilis MinD differs from the dimeric form by constructing MinD variants that are fixed in one of these conformations. Dimerization of MinD and other members of the MinD/ParA protein family requires binding of ATP. The conserved lysine in the Walker A-type ATPase domain, amino acid position 16 in the protein (Fig. S1), forms hydrogen bonds with the phosphate groups of the nucleotide, as revealed in the crystal structure of Pyrococcus furiosus MinD (39). Exchanging this residue into an alanine (K16A, apo monomer) prevents ATP binding and formation of dimers in several Walker motif-containing proteins, including E. coli MinD and RecA, and B. subtilis ParA/Soj (40–42). The conserved glycine at position 12 makes contacts with the γ-phosphate of ATP across the dimer interface (40). It has been shown that changing this residue into a valine in B. subtilis ParA causes a steric clash in the active site (G12V, ATP-bound monomer), thereby preventing dimerization while retaining ATP binding (40, 42). These two monomer-forcing amino acid exchanges (K16A and G12V) were introduced into B. subtilis MinD, which was fused at the N-terminus with GFP to follow its localization in the cell. It has been shown that a GFP-MinD fusion retains its biological activity (14), and the fusion protein was expressed at levels that resulted in normal sized cells and the absence of minicells (Fig. 1A). To prevent possible localization artefacts due to the weak dimerization characteristics of GFP (43), we used a monomeric GFP (mGFP) variant. The different MinD fusion proteins were expressed from the ectopic amyE locus, under control of the xylose-inducible Pxyl promoter (44), in a strain lacking wild type minD (ΔminD), as well as in a strain lacking minD and minC (ΔminCD). Western blot analysis showed that the different MinD-variants were stable and expressed at levels comparable to that of wild type MinD (Fig. S2). Induction of mGFP-MinD with 0.1 % xylose reduced minicell formation in a ΔminD background to wild type levels (from 14.7 % to 0.07 %, n > 300). However, the mGFP-MinD-K16A and -G12V variants were unable to prevent formation of minicells (11.2 % and 9.5 %, respectively, n > 600).

**Fig. 1.**
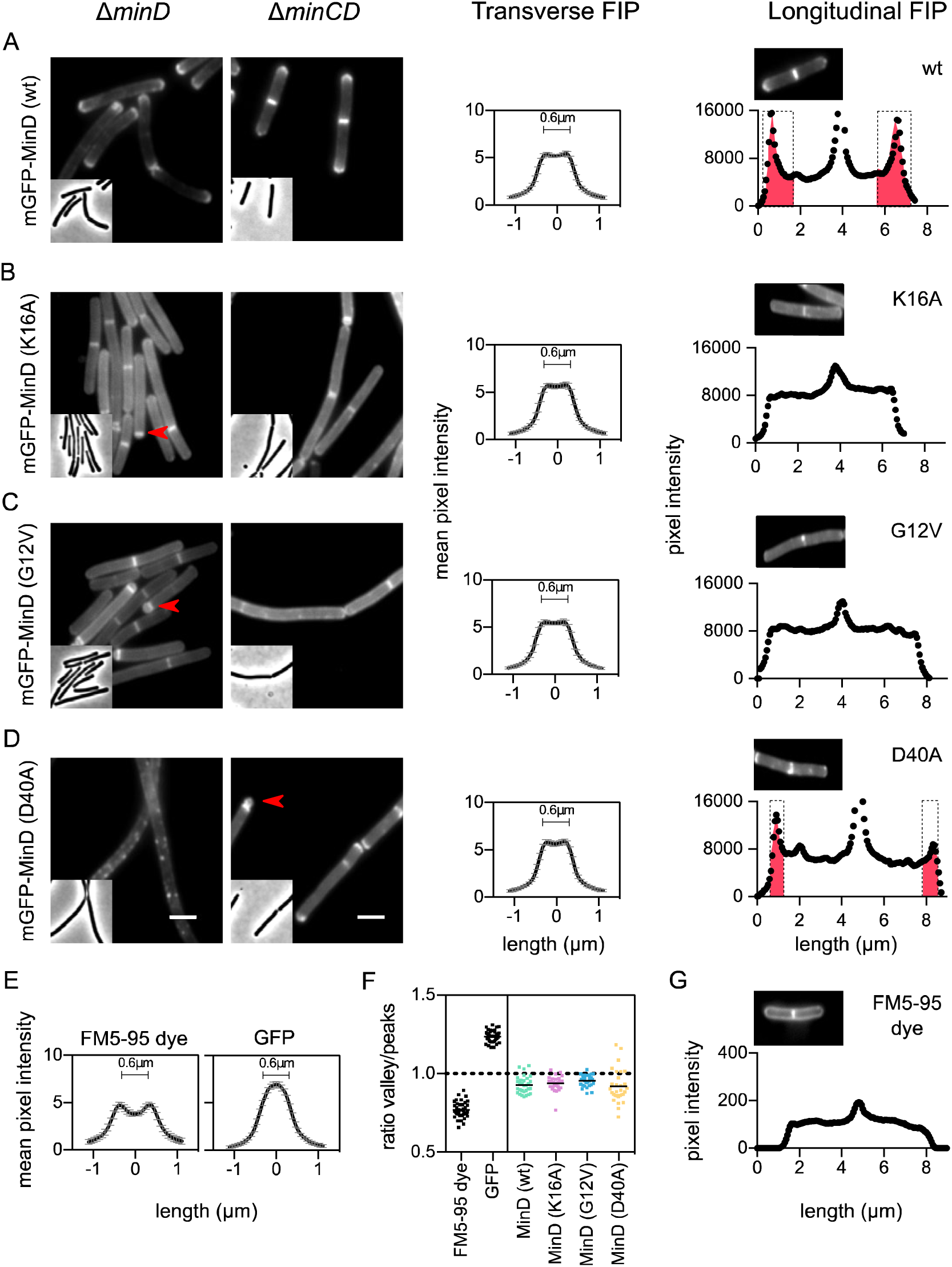
Localization of monomeric and dimeric MinD variants. Cellular localization of (A) wild type MinD, (B) MinD K16A, (C) MinD G12V, and (D) MinD D40A. Localization was monitored by an N-terminal mGFP fusion. Fusion proteins were expressed in either a ΔminD or ΔminCD background. Fluorescence images (left panels) and corresponding phase contrast images (inset) are shown in the left panels. Some minicells are indicated with red arrows. Scale bar is 2 μm. Middle panels show the transverse fluorescence intensity profiles (FIP) with standard deviations calculated using an average of at least 30 cells (ΔminCD background) per data set. Right panels depict the longitudinal fluorescence intensity profiles (FIP) using the ΔminCD background. The manually shaded red areas highlight the polar gradients. Additional examples for wild type MinD and the D40A variants are shown in Fig. S3. (E) Transversal fluorescence intensity profile (FIP) with standard deviations of exponentially growing wild-type cells stained with fluorescence membrane dye FM5-95, and wild-type cells expressing GFP are shown as controls. (F) Membrane affinities, with median values, estimated from the valley/peak ratios shown in the middle panels of (A-D) and controls (E). (G) Longitudinal fluorescence intensity profile (FIP) along the exponentially growing wild-type cells stained with the fluorescence membrane dye FM5-95. Strains used in (A): LB249 and LB305, (B): LB250 and LB306, (C): LB251 and LB307, (D): LB252 and LB308, (E): LB609 and (G) 168.

Fig. 1B shows the effect of the K16A exchange on mGFP-MinD localization. In line with previous results (38), preventing the binding of ATP abolishes the mGFP-MinD concentration gradient, which becomes clearer in longitudinal fluorescence intensity profiles (Fig. 1, right panels). The fluorescence peaks visible at midcell are caused by the presence of two septal cell membranes layers, as indicated by the fluorescence intensity profiles of a wild type cell stained with the fluorescent membrane dye FM 5-95 (Fig. 1G).

The monomeric ATP binding G12V variant shows the same absence of a protein gradient as the K16A variant (Fig. 1C). In contrast to E. coli MinD (29), both monomeric B. subtilis MinD mutants still bind to the cell membrane. To measure the membrane association in vivo, transverse fluorescence intensity profiles were collected (Fig. 1, middle panels), and the membrane affinities were estimated based on the ratio between valley (cytoplasmic) and peak (membrane) signals. This confirmed that the different monomeric mutants have a comparable membrane affinity as wild type MinD (Fig. 1F). As controls we used the fluorescence membrane dye FM5-95 and cells expressing cytosolic GFP (Fig. 1E).

It has been shown that exchanging a conserved aspartic acid residue in the ATPase domains of B. subtilis ParA and E. coli MinD blocks ATP hydrolysis and traps the proteins in a dimeric state (40, 42). When this aspartic acid residue, located at amino acid position 40 in the B. subtilis MinD sequence (Fig. S1), was replaced by an alanine (D40A, ATP-bound dimer, ATP hydrolysis deficient) in the mGFP-MinD fusion, expression of this variant in a ΔminD strain resulted in highly filamentous cells with clusters of mGFP-MinD-D40A formed along the membrane (Fig. 1D, left picture). Introduction of a minC deletion restored cell division (Fig. 1D, right picture), indicating that the locked MinD dimer causes hyper activation of MinC. Interestingly, in this ΔminCD background the clusters disappeared and mGFP-MinD-D40A showed a clear polar and septal accumulation, although the polar and septal signal intensities were approximately 60-70 % compared to that of wild type cells (Fig. S3B), and the longitudinal protein gradient was clearly reduced (Fig. 1D and S3A). Quantification of the membrane affinity showed that the membrane affinity of the D40A variant is only slightly higher compared to the monomeric mutants (Fig. 1F).

### Localization of trapped MinD dimer depends on MinJ

The polar and septal localization of the D40A trapped dimer-variant of MinD presumably relies on the interaction with MinJ. It has been reported that the absence of MinJ results in detachment of MinD from the membrane (32, 37). However, when we introduced the ΔminJ in our ΔminCD deletion strain background we found that all MinD-GFP variants remained attached to the membrane, and the septal and polar localization of wild type and the trapped dimer MinD disappeared (Fig. 2). These results show that, while MinJ is not required for membrane attachment of MinD per se, it is required for polar and septal localization of MinD through interactions specifically with the dimeric form of MinD.

**Fig. 2.**
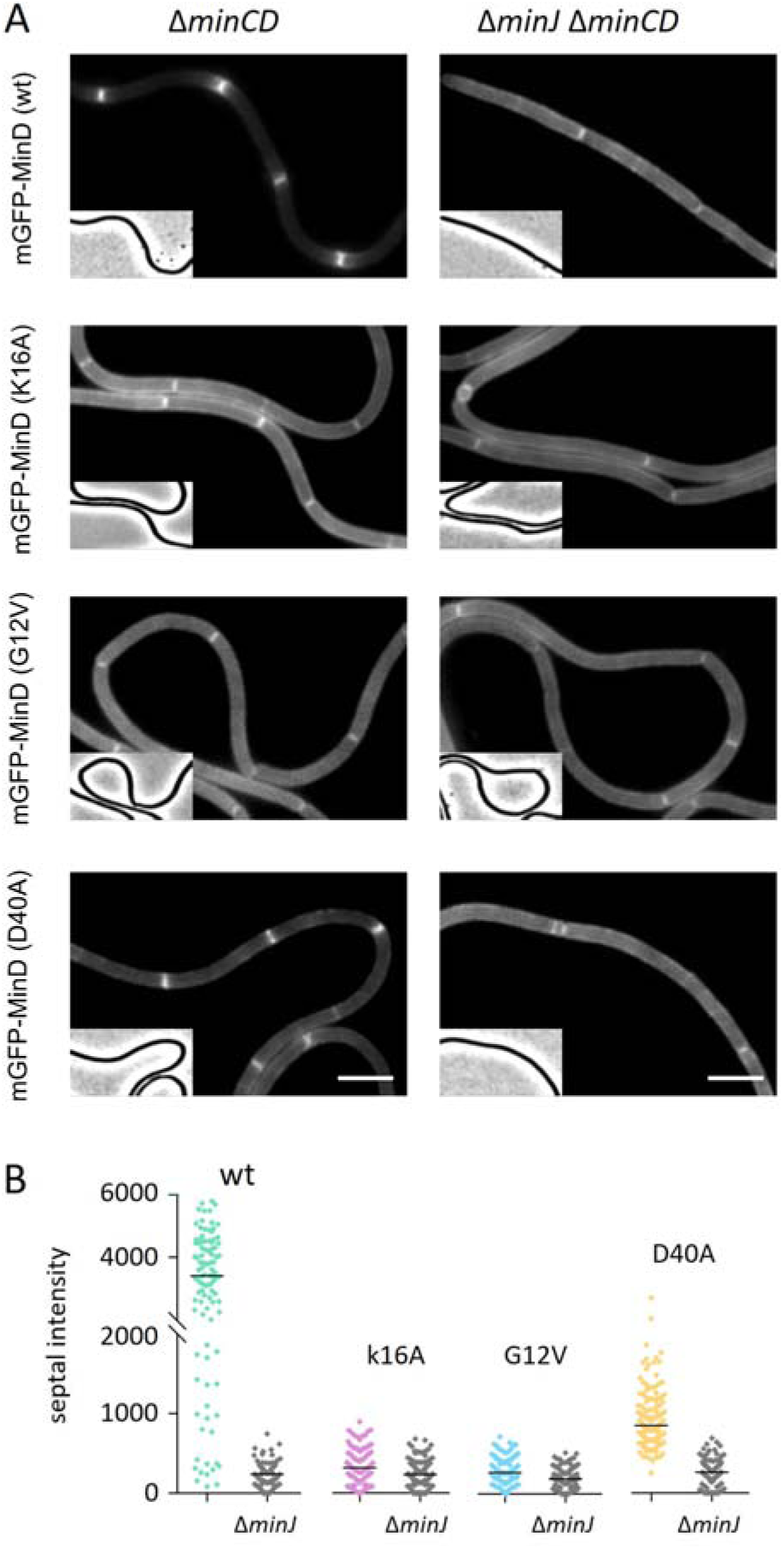
Effect of a minJ deletion on MinD-GFP localization. (A) Fluorescence microscopy images of ΔminCD and ΔminJ ΔminCD mutant cells expressing different mGFP–MinD variants. Corresponding phase contrast images are shown in the insets. (B) Quantification of related septal fluorescence intensities, with median values, at septa (n > 100). Since the strongly filamentous ΔminJ strain is delicate to handle, cells were grown on agarose patches on microscopy slides. Scale bar is 5 μm. Strains used for wt: LB405 and LB409, K16A: LB406 and LB410, G12V: LB407 and LB411, and D40A: LB408 and LB412.

### Recruitment of MinC by the different MinD mutants

Several biochemical studies with different bacterial species, including B. subtilis, have shown that MinC and MinD dimers can assemble into long polymers (45–48). However, the assembly of large MinCD multimers has not been observed in vivo, and a genetic study indicated that the formation of such polymers are not necessary for the function of the Min system in E. coli (49). The D40A MinD variant showed a clustered membrane localization pattern that resolved into a smooth membrane pattern when minC was deleted (Fig. 1D). This change in localization pattern might be related to the formation of higher-order MinCD assemblies. To confirm this, we introduced an active mCherry-MinC fusion to see whether mGFP-MinD and mCherry-MinC would colocalize when the former is fixed in a dimeric state. First, we examined the localization of MinC in the presence of monomeric K16A and G12V MinD variants. As shown in Fig. 3B and C, the mCherry-MinC signal was dispersed throughout the cytosol in these strains, indicating a lack of interaction with monomeric MinD variants, in line with a previous report (38). In case of the mGFP-MinD-D40A variant, the mCherry-MinC expression was only induced for 1 h to prevent excessive filamentation. As shown in Fig. 3D, the mCherry-MinC signal showed foci along the membrane that often colocalized with mGFP-MinD-D40A foci, confirming that MinD dimerization is crucial for MinC interaction, and that both proteins can indeed form large assemblies in the cell, presumably in the form of MinCD copolymers.

**Fig. 3.**
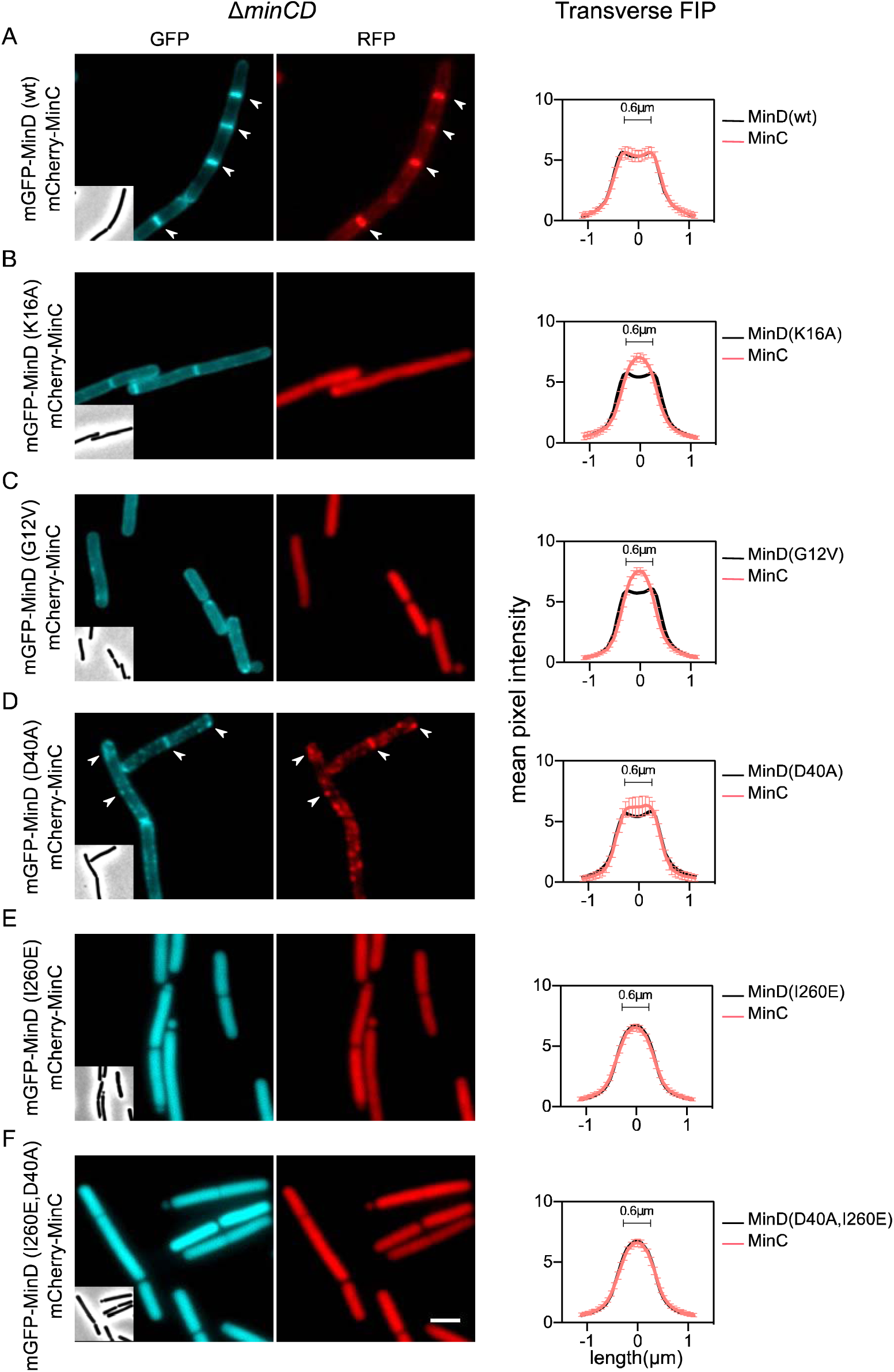
Membrane recruitment of MinC by MinD variants. Fluorescence microscopy images of cells expressing different mGFP–MinD variants (cyan) and mCherry–MinC (red). Corresponding phase contrast images shown in the insets. (A) Wild type MinD, (B) MinD K16A, (C) MinD G12V, (D) MinD D40A, (E) MinD I260E, (F) MinD D40A, I260E. Right panels show the transverse fluorescence intensity profiles (FIP) with standard deviations averaged over at least 30 cells. White arrows in (D) highlights colocalization. Scale bar is 2 μm. mGFP–MinD variants and mCherry–MinC were expressed in a ΔminCD background strain Strains used: (A) LB318, (B) LB319, (C) LB320, (D) LB321. (E) LB643 and (F) LB644.

The colocalization of the MinD-D40A and MinC fluorescent fusion proteins enabled us to investigate whether MinD dimerization is sufficient for MinC activity or whether localization at the cell membrane is also required. To test this, the membrane binding capacity of MinD was impaired by replacing isoleucine 260, located in the C-terminal membrane targeting amphipathic alpha helix, with a glutamate residue (Fig. 3E) (50). Next, we introduced this mutation in the mGFP-MinD-D40A mutant to create a trapped MinD dimer that was no longer able to bind to the membrane. Expression of this cytosolic MinD variant did not result in cell filamentation (Fig. 3F), indicating that interaction with MinD dimers alone is not sufficient to inhibit FtsZ polymerization. Rather, MinC needs to be localized and enriched to the membrane surface to be effective, which is not surprising since Z-ring formation occurs at the membrane periphery.

### Weak membrane interaction is important for MinD gradient formation

The absence of a clear difference in membrane affinity between monomeric and dimeric MinD raised the question whether a reversible membrane binding is important for the formation a protein gradient. To examine this, we tried to increase the membrane affinity of MinD by adding an extra copy of its C-terminal membrane targeting amphipathic helix. The extra amphipathic helix was connected to the original helix by a short 11 amino acid long flexible linker (Fig. 4A). First, we tested whether such tandem amphipathic helix (2xAH) increases the membrane affinity by fusing it to mGFP. As shown in Fig. 4, fusion of this tandem helix to mGFP increased the membrane association of the latter considerably when compared to a single MinD amphipathic helix.

**Fig. 4.**
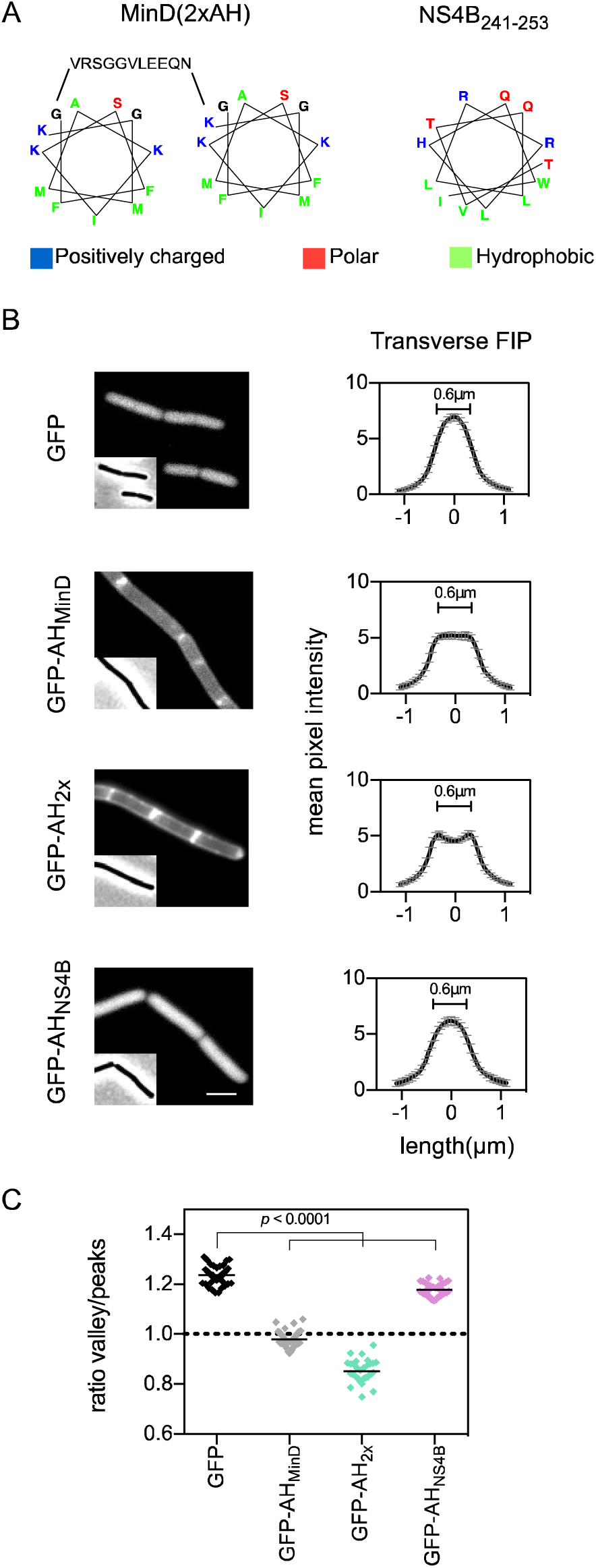
Membrane association of different amphipathic helices. (A) Schematic presentation of the tandem amphipathic helix and the weak amphipathic helix from Hepatitis C virus protein NS4B_241-253_. (B) Fluorescence microscopy images and transverse fluorescence intensity profiles (FIP) with standard deviations of cells expressing the different amphipathic helix sequences fused to the C-terminus of GFP. A strain expressing cytoplasmic GFP was included for comparison. Scale bar is 2 μm. Strains used: FBB043 (GFP-AH_MinD_), FBB05 (GFP-AH_2x_), FBB046 (GFP-AH_NS4B_), LB609 (GFP). (C) Average membrane affinities calculated from the transverse fluorescence intensity profiles (n > 30). Significance of difference was confirmed using t-test.

Next, we replaced the membrane targeting amphipathic helix of mGFP-MinD by the tandem amphipathic helix. Western blot analysis showed that this new C-terminus did not affect the protein levels to a significant degree (Fig. S2). As shown in Fig. 5B and D, the tandem helix notably increased the fluorescent membrane signal all along the cell membrane, thereby reducing the longitudinal fluorescence gradients.

**Fig. 5.**
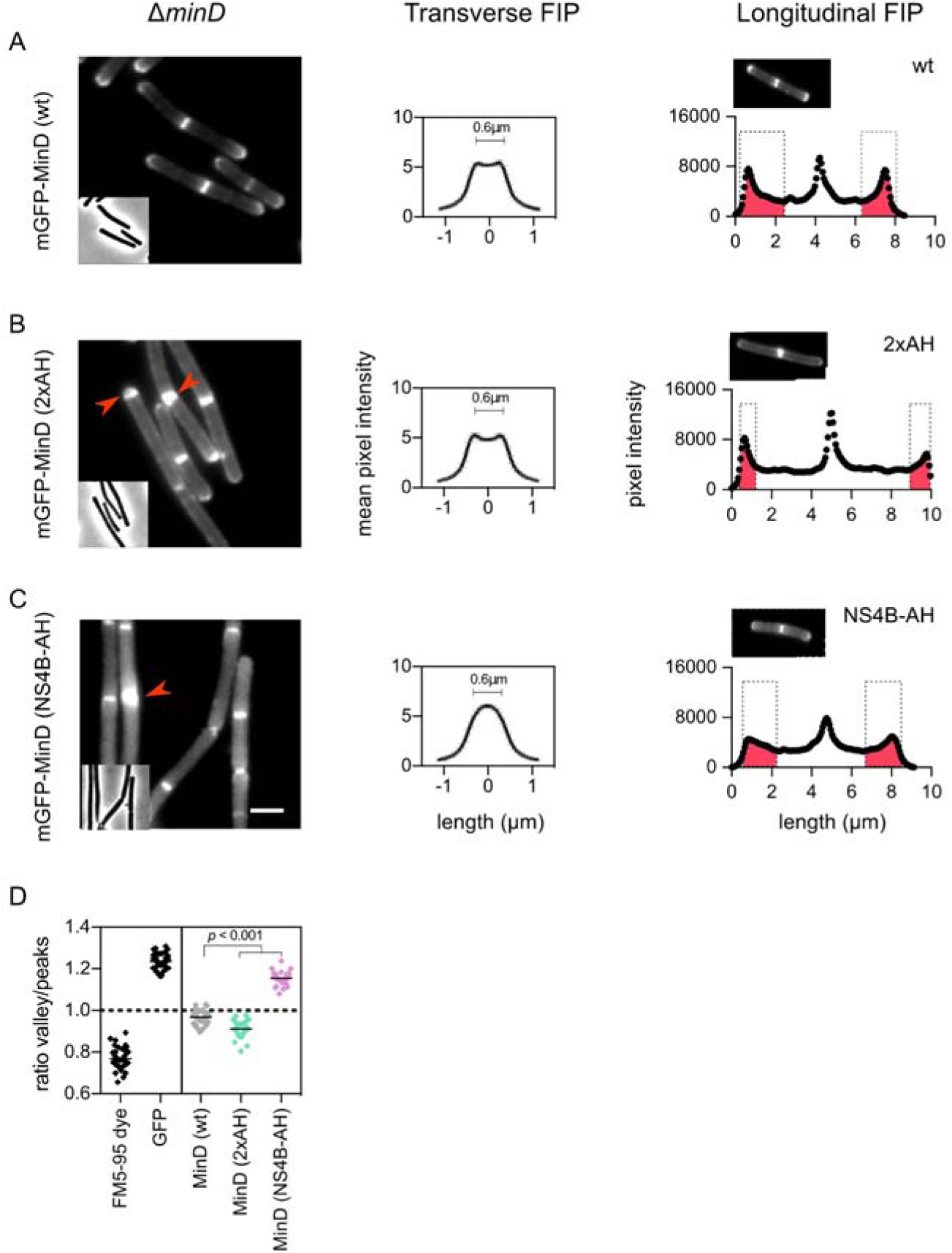
Membrane affinity affects MinD gradient. Fluorescence microscopy and fluorescence intensity profiles (FIP) with standard deviations of cells expressing mGFP-MinD containing either the native amphipathic helix (wt) (A), the tandem MinD amphipathic helix (2xAH) (B) or the weak Hepatitis C virus protein NS4B amphipathic helix (NS4B-AH) (C). The manually shaded red areas highlight the polar gradients. Additional examples are shown in Fig. S4. (D) Relative membrane affinities with median values of the different mGFP-MinD variants calculated from transverse fluorescence intensity profiles (n > 30). Significance of difference was confirmed using t-test. The MinD variants were expressed in a ΔminD background. Phase contrast images are shown as insets. Some double septa are indicated with red arrows. Scale bar is 2 μm. Red areas in the intensity profiles highlight the polar gradients. Strains used in (A) LB249, in (B) LB507, in (C) LB508.

To confirm that the tandem amphipathic helix increased the membrane affinity of MinD, we tested its sensitivity to membrane depolarization. Previously, it has been shown that binding of the MinD amphipathic helix to the cell membrane requires the presence of the membrane potential (50). Dissipation of the proton motive force by the specific proton-ionophore carbonyl cyanide m-chlorophenylhydrazone (CCCP) did not affect the membrane localization of the tandem amphipathic helix fusion, whereas CCCP quickly disturbed the membrane attachment of wild type MinD (Fig. S5).

We also tested what happens with the MinD gradient when its membrane affinity is reduced. For this, the well-studied membrane binding amphipathic helix 2 from the Hepatitis C virus protein NS4B was chosen (51, 52), which is hardly able to attach mGFP to the cell membrane (Fig. 4). Replacing the wild type membrane targeting helix of MinD by the weak NS4B amphipathic helix reduced the membrane association of mGFP-MinD (Fig. 5D), as well as its septal and polar localization (Fig. 5C and Fig. S4). Interestingly, this mutant still displayed a longitudinal protein gradient. Together these data suggests that a reversible membrane association is important for the formation of a MinD gradient along the cell axis.

### Effect of MinD membrane affinity on MinC recruitment and activity

The results shown in Fig. 3 indicated that the attachment of the MinD dimer to the membrane is essential for the activity of MinC. When this attachment was strengthened by using the tandem amphipathic helix, the respective strain regularly featured additional septa resulting in minicells (Fig. 5B, red arrows), although the frequency was lower compared to a ΔminD mutant (Fig. 6A). The strain also exhibited an elongated cell phenotype that was comparable to that of the minD deletion strain (Fig. 6B). As shown in Fig. 7B, the mGFP-MinD(2xAH) variant is capable of recruiting mCherry-MinC along the whole cell membrane, which likely explains the elongated cell phenotype.

**Fig. 6.**
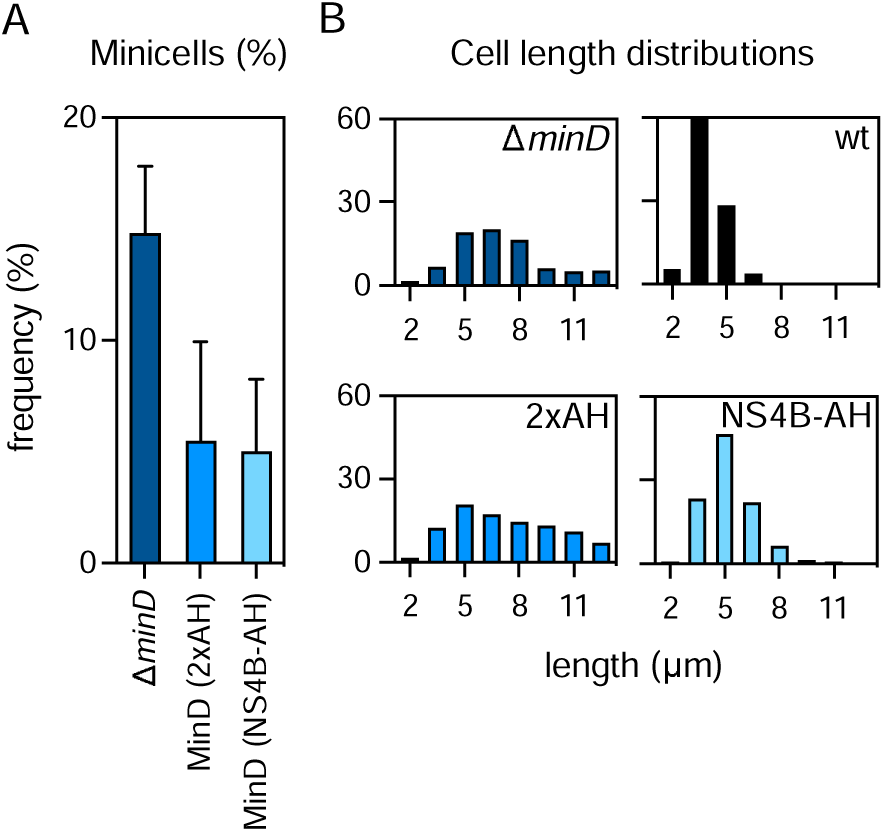
Functionality of MinD with different membrane affinities. (A) Minicell formation in cells expressing mGFP-MinD with different membrane affinities. The MinD variants were expressed in a ΔminD background (n > 300). Standard deviations are indicated. (B) Cell length distributions of the different strains (n > 300). Strains used: 1901 (ΔminD), LB249 (wild type MinD), LB507 (2xAH) and LB508 (NS4B-AH).

**Fig. 7.**
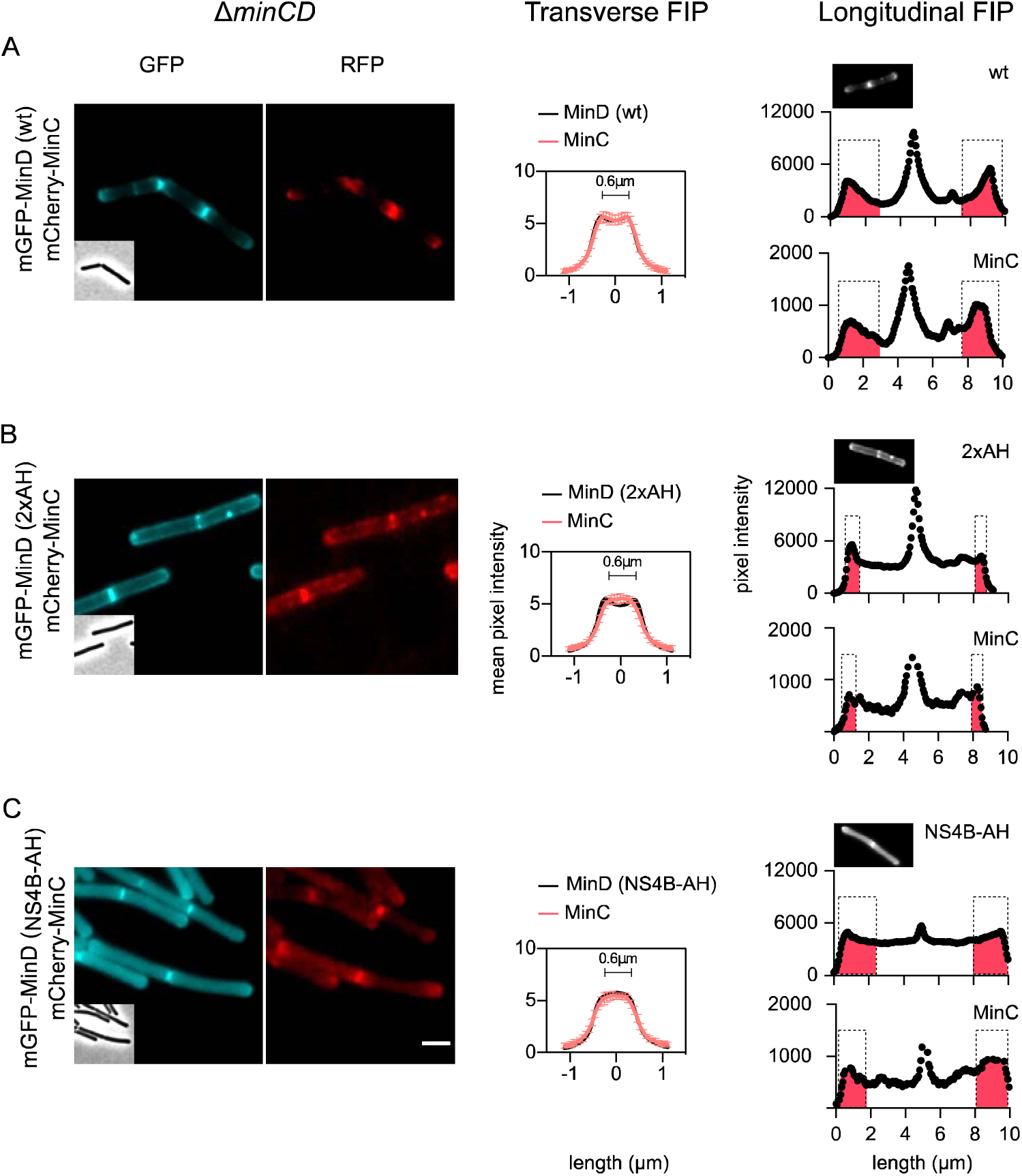
Increased MinD membrane affinity affects MinC recruitment. Fluorescence microscopy images of cells expressing mGFP-MinD (cyan) and mCherry-MinC (red), with either the native membrane anchor (A), the tandem amphipathic helix (B), or the weak Hepatitis C virus protein NS4B derived amphipathic helix (C). Corresponding phase contrast images are shown in insets. Scale bar is 2 μm. Transverse fluorescence intensity profiles (FIP) with standard deviations are shown in the right panels (n > 30). Right panels depict the longitudinal fluorescence intensity profiles (FIP) with manually shaded red areas to highlight the polar gradients. Additional examples are shown in Fig. S6. The mGFP-MinD variants and mCherry-MinC were expressed in a ΔminCD background strain. Strains used: (A) LB318, (B) LB584, (C) LB559.

Weakening the membrane affinity of mGFP-MinD, by replacing the wild type amphipathic helix with the NS4B amphipathic helix, also resulted in occasional minicell formation, but again not as numerous as in a minD deletion strain (Fig. 6A). The cell length distribution of the strain expressing the mGFP-MinD(NS4B-AH) variant was closer the wild type strain (Fig. 6B). When mCherry-MinC was co-expressed in this strain background, also the MinC fusion protein showed a clear fluorescent gradient along the cell axis (Fig. 7C). Thus, a reversible association of MinD with the cell membrane seems to be critical to prevent misplacing the FtsZ inhibiting activity of MinC along the cell axis. Both the mGFP-MinD(2xAH) and mGFP-MinD(NS4B-AH) variants were, at least partially, able to prevent minicell formation, presumably because they are still recruited to cell poles and cell division sites through interaction with MinJ.

### Membrane gradient formation in kinetic Monte Carlo simulations

The different experiments indicated that the formation of a MinCD gradient requires the dynamic switching between monomer and dimer states of MinD (Fig. 1), and a reversible MinD membrane interaction (Fig. 5). So far, in silico simulations of the B. subtilis Min gradient have assumed a strong difference in membrane interaction between the monomeric and dimeric state of MinD, which is based on properties of the E. coli Min system (36, 37). However, our data suggests that for B. subtilis these affinities do not differ much (Fig. 1). To investigate more closely whether a difference in membrane affinity between monomer and dimer is required for the formation of a MinD gradient in B. subtilis, we resorted to kinetic Monte Carlo simulations (see Methods for more details). The Monte Carlo algorithm attempts to move one molecule at a time by a small amount in a three-dimensional space. The new interaction energy experienced by the particle is then calculated and compared with its old energy, and the move is subsequently accepted or rejected according to the standard Metropolis test (53, 54). This procedure is used to guide the system to the correct Boltzmann distribution in equilibrium. It has been shown that this Monte Carlo dynamics corresponds well with Brownian or molecular dynamics (55, 56), and can be used to examine mechanisms underlying spatiotemporal protein gradients in cells (34), provided that the moves are small enough and that the acceptance probability remains high (55).

MinD monomers were presented as spheres (hydrodynamic radius 2.5 nm) that diffuse within and along a spherocylindrical cell geometry. The following rules were implemented: (i) Diffusion along the membrane is 10-fold slower that through the cytoplasm, and (ii) the dimer, due to its larger diameter, diffuses 2-fold slower compared to the monomer in both environments, emanating from Stokes-Einstein law and a hydrodynamic radius for the dimer which is double that of the monomer. (iii) Based on the data from Fig. 3, we started out with a 10 % stronger membrane affinity for the dimer. (iv) The membrane dwell time of monomers and dimers is 0.01 - 0.5 sec and during the simulations approximately 70 % of MinD molecules reside at the membrane, approaching the valley to peak values observed experimentally (Fig. 1F). (v) Dimerization and monomerization are modelled as chemical reactions between a monomer and a dimer state. Baseline rates are set such that the two states are in a 1:1 ratio, based on the longitudinal fluorescence intensity profile in Fig. 1A, and the assumption that dimers are located at the polar regions and monomers along the lateral wall. In our baseline model, dimerization at the pole, membrane and in the cytoplasm is equally likely, and is modelled in a simple way as a reaction turning a monomeric to a dimeric MinD at constant rate. (vi) The transition from dimer to monomer, stimulated by ATP hydrolysis, occurs stochastically with a half-life of approximately 1/sec, which is based on information from E. coli MinD (5). (vii) MinD dimers that come in close proximity of MinJ, represented as the surface area at the polar caps, will remain attached to MinJ for some time, so that approximately 25 % of MinD dimers is associated with MinJ, reminiscent of the septal localization of the D40A variant (Fig. S3). We simulated 10^7^ time steps, corresponding to about 100 s real time, when the system is in steady state. Simulation details are described in the material and methods, and a full list of parameters and their typical values considered in simulations is presented in Table S1.

When we applied these rules and ran the simulation, a strong accumulation of MinD at the poles was observed, but no gradient (Fig. 8A). In this simulation, the chance that MinD can form a dimer is the same for monomers in the cytoplasm and those that are attached to the membrane. However, it is likely that dimer formation in the latter situation is more likely since monomers attached to the membrane by their C-terminal amphipathic helix will diffuse slower and therefore spend more time in close proximity and, importantly, they have the same spatial orientation, which will facilitate dimerization. However, when we repeated the simulation, and implemented that membrane attached monomers have a 2-fold higher chance of forming dimers than monomers in the cytoplasm, the localization pattern did not change (Fig. 8B).

**Fig. 8.**
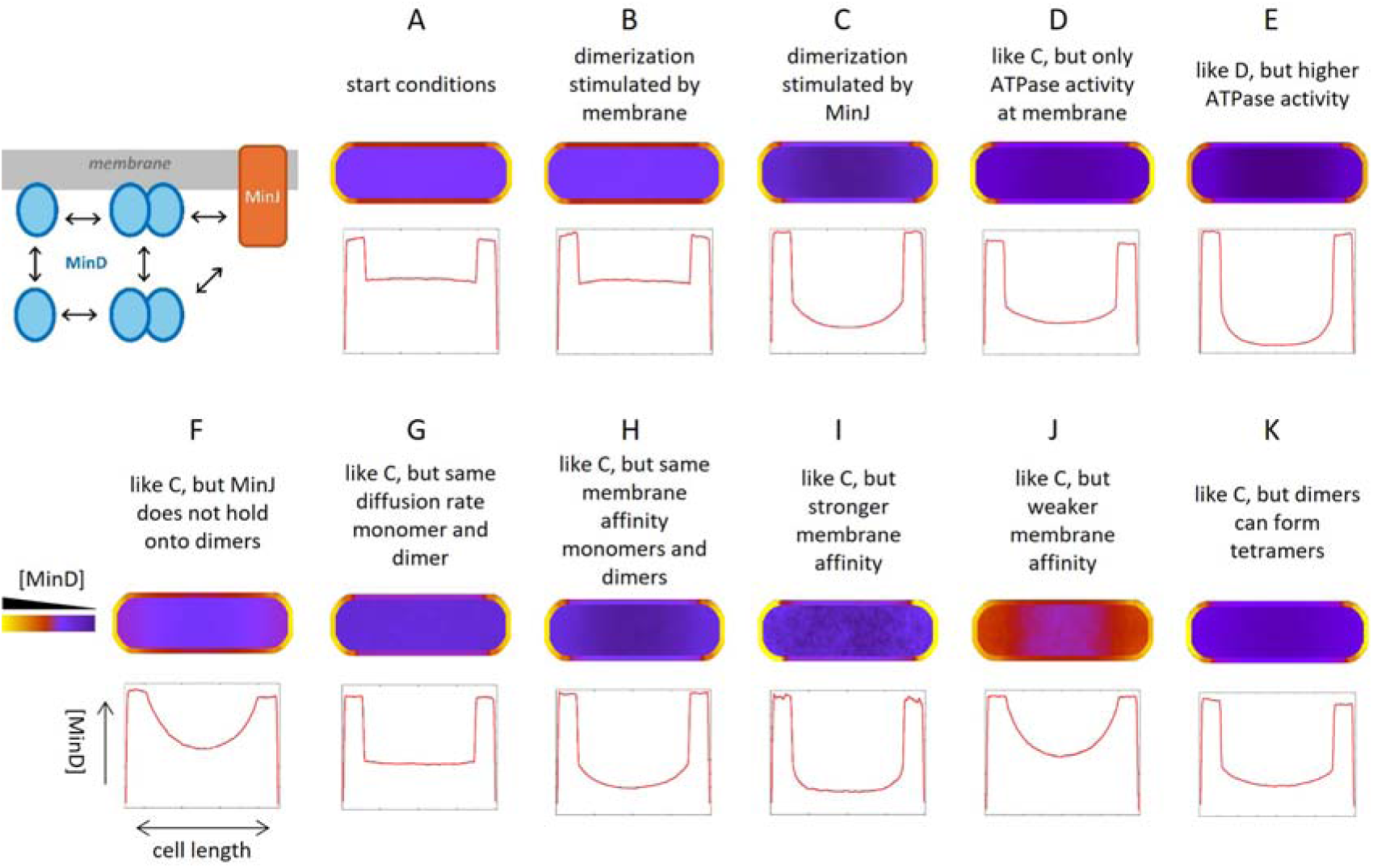
Kinetic Monte Carlo simulations of MinD localization. Whole-cell kinetic Monte-Carlo simulations of MinD distribution, taking into account dimer-to-monomer transition rates (ATPase activity), membrane affinities, and MinJ interaction (schematic model). The following conditions have been simulated: (A) Start situation whereby (i) diffusion along the membrane is 10-fold slower compared to cytoplasm, (ii) MinD dimer diffuses 2-fold slower compared to the monomer in both environments, (iii) 10 % stronger membrane affinity for dimer, (iv) membrane dwell time of monomers and dimers on average 1.4-4.5 sec, (v) dimerization and monomerization rates such that dimers and monomers are approximately in a 1:1 ratio, (vi) transition from dimer to monomer occurs stochastically with a half-life of approximately 1/sec, and (vii) MinD dimers in close proximity of polar regions (peak MinJ concentration) will remain attached for some time, so that approximately 25 % of MinD dimers is associated with the polar caps, representing MinJ. (B) Same as simulation A, but membrane attached monomers have a 2-fold higher chance of forming dimers compared to cytoplasmic monomers. (C) Same as simulation A, but MinJ also stimulates MinD dimerization. (D) Same as simulation C, but MinD ATPase activity, i.e. dimer-to-monomer transition, only occurs at the membrane. (E) Same as simulation D, but with a 10-fold higher ATPase activity, i.e. dimer-to-monomer transition. (F) Same as simulation C. but MinD dimers are not retained by MinJ. (G) Same as simulation C. but diffusion rates of monomer and dimer are the same. (H) Same as simulation C. but the membrane affinity of MinD monomers and dimers is the same. (I) Same as simulation C. but membrane affinity of MinD is stronger, such that the diffusion is a further 10-fold slower on the membrane. (J) Same as simulation C, but membrane affinity of MinD is 2-fold weaker. (K) Same as simulation C, but now MinD dimers can also form tetramers. Graphs indicate the average lateral projection of MinD in simulated cells.

Since it seems that MinJ interacts with the dimer form of MinD (Fig. 2), there is the possibility that MinJ actually stimulates MinD dimerization. Interestingly, when this property was added to the simulation, a clear MinD gradient emerged along the cell length, together with a strong polar localization signal (Fig. 8C), a pattern that is in good agreement with the longitudinal fluorescence intensity profiles observed in vivo (Fig. 1A). This pattern did not change when we included the condition that only membrane attached dimers are able to revert to monomers (Fig. 8D, E), in agreement with a recent report showing that the ATPase activity of MinD depends on its interaction with lipid membranes (57). In the simulation of Fig. 8E, we repeated the simulation of Fig. 8D, but increased the ATPase activity, i.e. dimer-to-monomer transition 10-fold. This enhances MinD gradient formation.

When we performed the simulation of Fig. 8C but now with the assumption that MinJ only stimulates dimerization and does not bind MinD dimers, a MinD gradient is still formed, although the strong accumulation at the polar caps was abolished (Fig. 8F). Subsequently, we repeated the simulation of Fig. 8C under conditions whereby there was no difference in the diffusion rate of monomer and dimer (Fig. 8G). In this case the MinD gradient was completely lost, showing that the differential diffusion of monomers and dimers is necessary to create a steady-state gradient.

Next, we tested the role of the membrane affinity in gradient formation. First, the simulation of Fig. 8C was repeated without a difference between the membrane affinity of the MinD monomer and dimer (Fig. 8H). Interestingly, under these conditions the MinD protein gradient is still being formed. When we repeated the simulation of Fig. 8C and kept the 10 % difference in membrane affinity between MinD monomer and dimer, but made their membrane affinities stronger, the gradient was reduced (Fig. 8I), in good agreement with our in vivo observation with a MinD variant containing a tandem membrane anchor (Fig. 5B). We then tested what happens when the simulation of Fig. 8C was repeated with a weaker membrane affinity of MinD. As shown in Fig. 8J, a clear gradient is formed, again in line with what was found when the membrane targeting amphipathic helix of MinD was replaced with the weaker amphipathic helix 2 from Hepatitis C virus protein NS4B (Fig. 5C).

It has been shown that purified E. coli MinD can form large polymers in the presence of ATP (58), and when bound to membranes can stimulate multimerization (59). Multimerization might further increase the diffusion difference between monomer and dimer, thus contributing to the formation of a protein gradient. Multimerization is challenging to model in our simulation setup, but as an approximation we tested what happens when MinD dimers can form tetramers. We ran a simulation whereby the baseline rates of dimer to tetramer and tetramer to dimer formations were equal. However this did not lead to a dramatic change in MinD gradient (Fig. 8K). In conclusion, our kinetic Monte Carlo simulations suggests that the formation of a MinD gradient requires: (i) a reversible and not too strong interaction of MinD with the membrane, (ii) a clear difference between the diffusion rates of monomer and dimer, (iii) an ATPase driven dimer-monomer cycle, and (iv) the stimulation of dimerization at the cell poles and division sites. Importantly, a difference in membrane affinity between monomer and dimer is not required.

## DISCUSSION

### Protein gradients in bacteria

Due to their small size, proteins diffuse rapidly throughout bacterial cells, in a matter of seconds, yet stable protein gradients can be found in a large variety of bacteria. What these systems have in common, including the MinD gradient, is a locally reduced diffusion rate of the proteins involved, which allows stable protein concentration gradients to form. The viscosity of the cell membrane is roughly 100 times larger compared to the cytoplasm (60, 61), and the Min system uses attachment to the cell membrane to slow down diffusion. Another approach to reduce diffusion is to bind to DNA, which occurs in the ParABS system involved in chromosome segregation in the asymmetric bacterium Caulobacter crescentus (62). This system consists of the ParA ATPase, which belongs to the same ATPase family as MinD (63) and ParB that binds to parS DNA binding sites. The ParA dimers associate with DNA nonspecifically whereby ParB stimulates ATP hydrolysis of ParA dimers, resulting in the dissociation from DNA. The localized dissociation and subsequent increased diffusion rate results in a ParA concentration gradient away from the parS locus (64). The cyanobacterium Synechococcus elongatus uses the ParA-like protein McdA to position carboxysomes along the longitudinal axis. In this system, the protein McdB, which interacts with carboxysomes, stimulates McdA ATPase activity and its release from DNA, resulting in an oscillation of McdA over the nucleoid (65). Recently it was described that Staphylococcus aureus trigger factor (TF) forms a gradient by interacting with the cell division protein FtsK, resulting in an increased TF concentration towards the septal region (66). How TF reduces its diffusion in S. aureus cells is still unclear but might be a consequence of the fact that the chaperone directly interacts with ribosomes (67), which diffuse much slower due to their size and association with mRNA and nascent polypeptide chains.

### Previous MinD gradient modelling studies

As mentioned in the introduction, there are many mathematical models describing the oscillating MinD gradient of E. coli but so far only two mathematical models focused on the MinD gradient of B. subtilis (36, 37). Both models make use of the reversible interaction of MinD with the cell membrane. The first study, almost 20 years ago, modelled the localization of DivIVA and MinD by a reaction-diffusion system in one spatial dimension (36). At that time neither MinJ nor the localization mechanism of DivIVA were known. In this early study both MinD and DivIVA were assumed to cycle between the membrane and the cytosol, and the membrane attachment of DivIVA would depend on MinD, whereas the membrane attachment of the latter would be stabilized at the cell poles by DivIVA. However, we now know that DivIVA binds to the membrane by itself, and there is no interaction between MinD and DivIVA (34).

The ATPase stimulated monomer-dimer cycle of MinD was not included in this early model, but it was used in a recent study that employed a minimal reaction-diffusion system in three-dimensional cellular space (37). In this model B. subtilis MinD cycles between the cell membrane and cytosol, whereby the membrane detached MinD is in the ADP-bound state and can only rebind after exchanging ADP for ATP, like in E. coli. By assuming that the membrane recruitment rate is higher at the poles and septa due to the presence of MinJ or due to a kinetic increase of diffusional impacts with the membrane due to the curved geometry at poles, a steep MinD gradient could be simulated (37).

### New data on MinD localization

Here, we have shown that monomeric MinD, both in its ATP-bound and unbound state, has a membrane affinity that is comparable to that of the dimeric form of MinD, which is clearly different from E. coli MinD that requires dimerization for membrane association. In fact, it has been shown that the C-terminal membrane targeting amphipathic helix of E. coli is considerably weaker than that of B. subtilis MinD, and that stable membrane binding is only possible when a dimer is formed so that there are two membrane targeting domains (24). Our findings are in good agreement with a recent report showing a strong membrane affinity for purified monomeric B. subtilis MinD for liposomes, independent of the presence of ATP (57).

We have also shown that MinJ is not required for the membrane recruitment of MinD (Fig. 2). We have no explanation for the differences between our findings and that of the previous studies in which it has been shown that MinJ is needed for the membrane localization of MinD (32, 37). However, since the membrane-targeting amphipathic alpha helix of B. subtilis MinD is by itself sufficient to recruit GFP to the cell membrane, even in E. coli, which does not contain a MinJ homologue (24, 50), it is difficult to envision how the absence of MinJ would bring MinD to the cytosol.

In another study, it has been suggested that the C-terminal amphipathic helix of MinD by itself is sufficient to recruit GFP to cell division sites, even in the absence of either MinJ or DivIVA (68). However, as indicated in Fig. 1, the presence of two perpendicular-observed membranes that are formed during septum synthesis gives a higher fluorescence membrane signal, which might have been mistakenly interpreted as septal localization. In addition, in our hands a GFP fusion with the C-terminal amphipathic helix of B. subtilis MinD shows a clear fluorescence membrane stain but no accumulation at division sites (Fig. 4B).

### Kinetic Monte Carlo model

Our in vivo findings warranted a new mathematical model to study the formation of the B. subtilis MinD gradient (see Methods for more details). The kinetic Monte Carlo model showed that a membrane attachment-detachment cycle is not required to form a protein gradient, but that a dynamic ATP-driven cycle between monomer and dimer is crucial, coupled to the differences in diffusion rates. We have shown that the MinD dimer associates with MinJ, but this by itself, even when the interaction was reversible, was not enough to create a gradient. We ascribe this to the fact that both monomers and dimers bind to the membrane with small difference in affinities in our model, whereas in (37) the monomer state is not membrane-associated. To establish a gradient in our simulations, we needed to include the requirement that MinJ stimulates and/or stabilizes MinD dimerization, but this MinJ activity remains speculative. It should be stressed that MinJ is not crucial for MinD dimerization, since a minJ mutant shows a strong filamentous phenotype that can be suppressed by a minC deletion, indicating that MinD dimerization, required for MinC activity, also occurs in the absence of MinJ.

Although a difference in membrane affinity between monomer and dimer is not necessary to establish a gradient, the modelling shows that a transient and reversible membrane binding of MinD is necessary. This was also confirmed by the reduced formation of a gradient when the membrane affinity of MinD was increased by using a tandem C-terminal amphipathic helix (Fig. 5B). This can be explained by considering the emergent length scale over which the gradient develops. As in reaction-diffusion models for morphogenesis, this length scale is given by the square root of a diffusion coefficient over a reaction rate. Our simulations suggest that a possible formula is given by 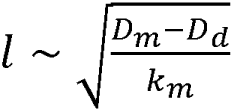, with *D_d_* and *D_m_* the dimer and monomer diffusion (the relevant rate is that on the membrane), and *k_m_* the rate of conversion between dimers and monomers. Increasing the interaction with the membrane leads to a decrease in diffusivity, and concomitantly a reduction in the length scale over which the gradient can be observed.

In our modelling, MinJ is simply represented by the polar cap area. FRAP studies have shown that the recruitment of MinJ to cell poles and division sites is quite dynamic with a fluorescence recovery rate close to a minute (37). However, the fluorescence recovery rate of MinD is approximate 8-fold higher (37). Because of this large difference in diffusion rates, we have ignored the diffusion of MinJ in our modelling. We do not foresee different results if we were to include MinJ diffusion in our Monte Carlo simulations, provided that in steady state MinJ is on average localized at the poles.

Another aspect that we have only partially addressed in the simulations is MinD polymerization. It has been shown that purified E. coli MinD can form large polymers in the presence of ATP (58), and, when bound to membranes, can stimulate multimerization (59). In our simulations we tested the effect of MinD tetramerization, however, this did not improve MinD gradient formation (Fig. 8K). It should be mentioned that formation of large polymers was not found with purified B. subtilis MinD (48). Moreover, FRAP experiments with B. subtilis shows a fluorescence recover rate of seconds for MinD at division sites (37), suggesting that if such polymers exist, they are not very stable. Thus, it remains to be seen whether MinD multimerization is relevant in B. subtilis cells.

In conclusion, we have shown here that a dynamic cyclic change in membrane affinity of MinD is not required for the formation of a concentration gradient in the cell. Instead, our modelling suggests that stimulation of MinD-dimerization at cell poles and cell division sites is needed. It will be interesting to see whether future experiments proves this suggestion right or wrong.

## MATERIAL AND METHODS

### Bacterial strains, growth conditions and media

Luria-Bertani (LB) agar was used for the routine selection and maintenance of both B. subtilis and E. coli strains. For B. subtilis, cells were grown in LB medium or Spizizen minimal salt medium supplemented with 0.05 % yeast extract (SMMY). SMMY consists of 2 g/l (NH_4_)_2_SO_4_, 14 g/l K_2_HPO_4_, 6 g/l KH_2_PO_4_, 1 g/l sodium citrate, 2 g/l MgSO_4_, 5 g/l fructose, 2 g/l tryptophan, 0.2 g/l casamino acids and 2.2 g/l ammonium ferric citrate. All strains were grown at 37 °C. The supplements were added as required: 100 µg/ml ampicillin, 5 µg/ml chloramphenicol, 5 µg/ml kanamycin, 100 µg/ml spectinomycin, 10 µg/ml tetracycline and 1 µg/ml erythromycin. The B. subtilis strains used are listed in Table S2. The mutant strains kindly provided by other labs were transformed into our laboratory strain to ensure that all strains were isogenic.

### Strain constructions

Molecular cloning, PCR and transformations were carried out by standard techniques. Plasmids and oligonucleotides used in this study are listed in Table S3 and S4, respectively. The minC, minD, minCD and minJ gene deletions were introduced from existing strains listed in Table S2 by standard natural transformation (69). The aprE-integration vector containing mCherry under an IPTG-inducible promoter (Pspac) was constructed as follows. The spectinomycin marker of pAPNC213 (70) was replaced with a chloramphenicol resistance cassette (cat) using In-Fusion cloning (Clonetech). For this aim pAPNC123 was PCR-linearized with oligonucleotide pair HS05/HS06 and the cat cassette was amplified from pSG2 (71) with oligonucleotide pair HS07/HS08. These PCR products were fused together with In-Fusion cloning, resulting in pAPNCcat, this plasmid was verified by sequencing. Next, the mCherry encoding gene was amplified using oligonucleotide pair HS437/HS438 and plasmid template pSS153 (72) as template. SalI and BamHI restriction sites were inserted into the primers. The mCherry PCR product and the pAPNCcat were digested with SalI and BamHI and ligated. The resulting plasmid was verified by sequencing and named pHJS112.

The IPTG-inducible mCherry-minC fusion was constructed as follows. A PCR fragment containing minC was amplified with oligonucleotide pair LB1/LB2 and genomic DNA of the wild type strain 168 as template. BamHI and EcoRI restriction sites were inserted into the primers. The PCR product and the aprE-integration vector pHJS112 were digested with BamHI and EcoRI restriction enzymes and ligated. The resulting plasmid pLB11 was verified by sequencing and transformed into B. subtilis competent cells, resulting in strain LB31. The aprE integration was verified by PCR, amplifying the genomic DNA with oligonucleotide pair HS508/HS509 and sequencing.

The amyE-integration vector pSG1730 containing GFP fused to minD under a xylose-inducible promoter (13) was used to exchange the minD residue glycine 12 to valine (G12V) and aspartic acid 40 to alanine (D40A), by incorporation of the corresponding mutation using the QuickChange site-directed mutagenesis reaction (Stratagene) and the oligonucleotide pairs HS320/HS321 and HS322/HS323, respectively. The resulting plasmids were verified by sequencing and named pHJS115 and pHJS116, respectively. The plasmid pHJS113 containing GFP fused to B. subtilis minD coding sequence with the lysine 16 to alanine (K16A) exchange under a xylose-inducible promoter was reported previously (50).

Plasmid pSG1729 (44), containing GFP under a xylose-inducible promoter, was used to generate a construct with GFP fused to the native minD membrane targeting amphipathic helix (KGMMAKIKSFFG). This domain was amplified from B. subtilis wild type strain 168 with oligonucleotide pair FB136/FB149. EcoRI and BamHI restriction sites were inserted into the primers. The PCR product and the plasmid pSG1729 were digested with EcoRI and BamHI restriction enzymes and ligated. The new plasmid was verified by sequencing and named pHJS117. Plasmid pHJS117 was used to generate a construct with GFP fused to two minD amphipathic helices in tandem with a short linker of glycines in between (2xAH). This tandem membrane targeting sequence was created by amplification with oligonucleotides FB138/FB149 and pHJS117 as template. The primer FB138 carries a long overhang that generates an extra MinD membrane targeting amphipathic helix fused to the existing membrane targeting domain (KGMMAKIKSFFGVRSGGVLEEQNKGMMAKIKSFFG). EcoRI and BamHI restriction sites were inserted into the primers. The PCR product and the plasmid pSG1729 were digested with EcoRI and BamHI restriction enzymes and ligated. The new plasmid was verified by sequencing and named pHJS119.

Another construct with GFP fused to the NS4B amphipathic helix 2 from Hepatitis C virus (NS4B-AH) was generated as follows. Plasmid pSG1729 was amplified with oligonucleotides FB141/FB149. The primer FB141 carries a long overhang that generates the NS4B-AH anchor (TVTQLLRRLHQWI). EcoRI and BamHI restriction sites were inserted into the primers. The PCR product and the plasmid pSG1729 were digested with EcoRI and BamHI restriction enzymes and ligated. The new plasmid was verified by sequencing and named pHJS121.

We constructed two different amyE-integration plasmids with GFP fused to minD with different membrane targeting sequences. The native minD amphipathic helix was either replaced with two minD amphipathic helices in tandem with a short linker of glycines in between (MinD2xAH) or with the Hepatitis C virus NS4B amphipathic helix 2 (MinDNS4B-AH). The minD gene was amplified from the B. subtilis genome (strain 168) with oligonucleotides pairs FB135/FB138 and FB135/FB141, respectively. BamHI and EcoRI restriction sites were inserted into the primers. The primers FB138 and FB141 carry a long overhang that generates the 2xAH and the NS4B-AH, respectively. The digested PCR fragments were ligated with BamHI/EcoRI-linearized pSG1729, containing GFP under a xylose-inducible promoter. The new plasmids were verified by sequencing and named pHJS123 and pHJS125, respectively.

Plasmids pSG1730, pHJS113, pHJS115, pHJS116, pHJS123 and pHJS125 were used as template for the QuickChange site-directed mutagenesis reaction (Stratagene) with oligonucleotide pair HS410/HS411 to introduce the A206K exchange in the GFP coding sequence to reduce protein dimerization (mGFP). The resulting plasmids were verified by sequencing and named pLB21, pLB22, pLB23, pLB24, pLB49 and pLB50 respectively. Plasmids pLB21 and pLB24 were used to further exchange the minD isoleucine 260 to glutamic acid (I260E) by QuickChange site-directed mutagenesis reaction (Stratagene) with oligonucleotide pair HS205/HS206. The resulting plasmids were verified by sequencing and named pLB71 and pLB72, respectively. These constructs were integrated at the amyE locus in different genetic backgrounds.

### Western blot analysis

To detect the cellular levels of the different GFP-MinD variants the strains were induced with 0.1 % xylose for three doublings. 2 ml of cultures, with comparable ODs were centrifuged and the cell pellets flash frozen in liquid nitrogen. The pellets were then resuspended in 200 µl of buffer containing 10 mM Tris-Hcl pH 8, 100 mM NaCl and 1 % TritonX-100. The cells were lysed with 100 µg/ml lysozyme during 15 min incubation at room temperature. Cell debris were spun down (20 min Eppendorf centrifuge, 14000 rpm, 4°C) and 10 μl of samples were loaded onto a 10 % SDS-PAGE followed by separation by electrophoresis at 150 V for 60 min. The proteins were then transferred to a PVDF membrane (Thermo Scientific) with transfer buffer (14.4 g/l glycine, 3 g/l Tris base and 15 % methanol) using mini-wet electroblotting system (Bio-Rad) at 55 V for 130 min. The transferred membrane was then blocked in blocking buffer (1 % skimmed milk powder in TBST buffer 50 mM Tris, 150 mM NaCl and 0.05 % Tween-20) overnight and rinsed twice for 5 min in TBST. The proteins were probed with a 1:2000 dilution of rabbit anti-GFP primary antibody (Invitrogen) in 10 ml TBST incubated 1 h, the membrane was washed 3x in TBST and incubated in a 1:20 000 dilution of secondary antibody goat anti rabbit IR dye (Li-cor Biosciences) in 10 ml TBST during 1 h. The immunoblot was developed by using an Odyssey® imaging system (Li-cor Biosciences).

### Microscopy

Microscopy was performed on an inverted fluorescence Nikon Eclipse Ti microscope equipped with a CFI Plan Apochromat DM 100x oil objective, an Intensilight HG 130bW lamp and C11440-22CU Hamamatsu ORCA camera. Images were analysed using ImageJ v.1.48d5 (National Institutes of Health) software and the plugin ObjectJ (73). To measure the transverse fluorescence intensity profiles (FIP), line-scans perpendicular to the cell axis were marked. The polar gradient areas, marked in red in figures, were manually assigned by means of visual inspection of the fluorescence gradient of the FIPs. The membrane affinity was estimated as the ratio valley/peaks of the transverse FIP, using an average of at least 30 cells per data set. When appropriate, data was analyzed with t-tests (p-values) using GraphPad Prism for Mac, GraphPad Software.

For microscopy, cells were grown to exponential phase by diluting 1:100 a starter culture grown overnight in liquid medium (SMMY) into fresh medium, and allowing at least three doublings before mounting on microscopy slides covered with a thin film of 1 % agarose. 0.1 % xylose and 0.01 mM IPTG were used for the induction since these concentrations complemented a ΔminCD deletion strain, preventing minicell formation. Membranes were stained with the fluorescent membrane dye FM-95 (0.5 µg/ml final concentration), incubated for 5 min prior to mounting onto microscope slides covered with a thin film of 1 % agarose. Dissipation of the proton motive force (pmf) was carried out by the addition of 100 μM carbonyl cyanide m-chlorophenylhydrazone (CCCP) for 10 min. CCCP was dissolved in DMSO (0.1 % final concentration of DMSO). As control, cells were incubated with 0.1 % DMSO.

### Kinetic Monte Carlo simulations

To model the formation of a steady state gradient in B. subtilis wild-type cells and mutants, kinetic Monte Carlo simulation were performed. Kinetic Monte Carlo is a stochastic algorithm which is known to be equivalent to molecular dynamics for sufficiently small local

moves (55, 56). The system simulated consisted in a collection of 1000 MinD proteins enclosed inside a spherocylinder, with cap diameter 2R_cyl_ and total height h_cyl_ (we chose yan aspect ratio 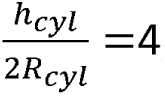, sizes 80 by 20 in simulation units) modelling a B. subtilis cell. MinD proteins can be in one of two states, either monomeric (M) or dimeric (D). The M and D states have both affinities for the membrane (the surface of the spherocylinder). To model this, we introduce an attractive potential, a square well with range 1 and strength *ε_M,_* and *ε_D_* for the M and D state respectively. Given our experimental results that the dimer is only weakly stickier than the monomer, we ran our simulations with *ε_D_* - 1.1 *ε_M_*, and *ε_M_*=1 k_B_T. Additionally, the D state, but not the M state, may interact with the caps (poles) in the spherocylinders, with an extra interaction again represented by a square well, with range 1 and strength *ε_C_*. This interaction simulates binding between dimeric MinD and MinJ. Various cases were considered but results are reported, for simplicity for two cases only: *ε_C_*=0 (no additional polar interaction), or *ε_C_*= 0.7 *ε_M_* . A third parameter which varies between M and D is the diffusion coefficient: for simplicity, unless otherwise stated, we assume this to scale inversely with molecular weight, so that the diffusivity of the dimer, *D_d_*, is half that of the monomer, *D_m_*.

Key to the formation of the gradient is the fact that the reaction between the M and D states of MinD is not in thermodynamic equilibrium, as it is linked to ATP hydrolysis. In our model, D turns into M, due to ATP hydrolysis, at a constant rate k_ATP_, in all cases. We consider instead three cases for the dimerization reaction. In our baseline model the conversion of M into D also occurred at the same rate in the cytoplasm, at the membrane and at the pole. We have set this rate such that the M and D state are on average comparable in steady state, as suggested by our experiments. We also considered the case where dimerization at the membrane occurs 2-fold more often at the membrane, and a third case where polar localization (through MinJ interaction) strongly stimulates dimerization, so that the M to D transition occurs at a rate k_pole_ >> k_ATP_ at poles. When considering polar stimulation, dimerization occurs at a rate k_ATP_/10 in the cytosol; otherwise, rates of dimerization and monomerization in the cytoplasm are equal. The diffusion of monomeric MinD was set to 0.0033 in simulation units (corresponding to approximately 2 micron square per second, see below), where the hydrolysis rate k_ATP_ was set to 10^-5^ (corresponding to 1/s, which is of the same order as used in previous modeling studies (5)). When considering polar stimulation, we set k_pole_ such that dimerization at the pole occurs with unit probability during a timestep (we have verified though that the results are qualitatively similar down to at least a value of k_pole_ equal to 100 k_ATP_). More details on the parameters used in the model are given in the caption of Fig. 8, and Table S1.

To simulate the monomer (K16A) and dimer (D40A) mutants, we used another version of the same model, in which all particles were of type M or D respectively, and the rate of interconversion between the two was set to 0.

Simulation units may be mapped to physical units by noting that 1 space and time units in simulations can be made to correspond to 0.08 microns and 10 microseconds, respectively. Simulations were run typically for 10^7^ time steps. Snapshots shown are for systems which have reached equilibrium, so representing steady-state behaviour.

## REFERENCES

## ACKNOWLEDGEMENTS

We would like to thank especially Norbert Vischer for analysis and quantification of fluorescence images and developing ObjectJ plugins, members of the Bacterial Cell Biology group (University of Amsterdam) for insightful discussions, Dirk-Jan Scheffers for critical reading of the manuscript, and Marc Bramkamp for prior communications. This research was funded by EU Marie Curie ITN grants ATP-BCT (020496–2) and AMBER (317338), Marie Curie CIG grant DIVANTI (618452) and STW Vici grant (12128).

