## Supplementary Information for "Membrane affinity difference between MinD monomer and dimer is not crucial for MinD gradient formation in Bacillus subtilis"

Table S1 Table with parameter values in simulation and physical units

Table S2 List of *B. subtilis* strains used in this study

Table S3 Plasmids used in this study

Table S4 Oligonucleotides used in this study

Fig. S1 Amino acid sequence alignment of *E. coli* and *B. subtilis* MinD proteins

Fig. S2 Western blot analysis of mGFP-MinD variants

Fig. S3 Longitudinal fluorescence intensity profiles of mGFP-MinD and D40A variants

Fig. S4 Longitudinal fluorescence intensity profiles of mGFP-MinD with different membrane anchors

Fig. S5 Effect of membrane potential dissipation on membrane binding

Fig. S6 Longitudinal fluorescence intensity profiles of mCherry-MinC in cells expressing different mGFP-MinD membrane-binding mutants.

References

**Table S1. Table with parameter values in simulation and physical units**

Mapping has been made considering one spatial and temporal simulation units correspond to $0.08\,\mu m$ and $10\,\mu s$ respectively; different assumptions will lead to different mapping which can be calculated by scaling the values shown in this Table according to the physical units of the corresponding quantity. As described in the Methods, we have called $D_m, D_d$ the diffusion coefficient of the monomer and the dimer respectively; whereas $p_{MD}^{pole}, p_{MD}^{mem}, p_{MD}$ denote the rates of monomer to dimer conversion at the pole, membrane and in the bulk respectively, and $k_m = k_{ATP}$ denotes the rate of dimer to monomer conversion. Other parameters are: (i) $\epsilon_M, \epsilon_D, \epsilon_C$, which are respectively the interaction energy between membrane and monomer, between membrane and dimer, and between spherocylinder cap and dimer; (ii) $R_{cyl}$ and $h_{cyl}$, which denote the radius and height of the spherocylinder that models the bacterium.

| Parameter | Value (Simulation Units) | Value (Physical Units) |
|---|---|---|
| $D_m$ | 0.0033 | $\simeq 2.1\,\mu m^2/s$ |
| $D_d$ | 0.0017 | $\simeq 1.1\,\mu m^2/s$ |
| $p_{MD}^{pole} = k_{pole}$ | 1 | $10^5 s^{-1}$ |
| $p_{MD}^{mem}$ | $2 \times 10^{-6}$ | $0.2\,s^{-1}$ |
| $p_{MD}$ | $10^{-6}$ | $0.1\,s^{-1}$ |
| $k_m = k_{ATP}$ | $10^{-5}$ | $1\,s^{-1}$ |
| $\epsilon_M$ | 1 | $k_B T \simeq 4.1 \times 10^{-21}$J |
| $\epsilon_D$ | 1.1 | $1.1\,k_B T$ |
| $\epsilon_C$ | 0.7 | $0.7\,k_B T$ |
| $R_{cyl}$ | 10 | $0.8\,\mu m$ |
| $h_{cyl}$ | 80 | $6.4\,\mu m$ |

**Table S2. List of *B. subtilis* strains used in this study**

The mutant strains kindly provided by other labs were transformed into our laboratory strain to ensure that all strains were isogenic.

| Strain | Genotype | Reference |
|---|---|---|
| 168 | wild-type | Lab stock (1) |
| 1901 | *minD::erm* | (2) |
| 3381 | *minC::km* | (3) |
| 3309 | *minCD::km* | (3) |
| RD021 | *minJ::tet* | (4) |
| LB249 | *minD::erm amyE::spc Pxyl-mGFP-minD* | This study |
| LB250 | *minD::erm amyE::spc Pxyl-mGFP-minD(K16A)* | This study |
| LB251 | *minD::erm amyE::spc Pxyl-mGFP-minD(G12V)* | This study |
| LB252 | *minD::erm amyE::spc Pxyl-mGFP-minD(D40A)* | This study |
| LB305 | *minCD::km amyE::spc Pxyl-mGFP-minD* | This study |
| LB306 | *minCD::km amyE::spc Pxyl-mGFP-minD(K16A)* | This study |
| LB307 | *minCD::km amyE::spc Pxyl-mGFP-minD(G12V)* | This study |
| LB308 | *minCD::km amyE::spc Pxyl-mGFP-minD(D40A)* | This study |
| LB318 | *minCD::km amyE::spc Pxyl-mGFP-minD aprE::cat Pspac-mCherry-minC* | This study |
| LB319 | *minCD::km amyE::spc Pxyl-mGFP-minD(K16A) aprE::cat Pspac-mCherry-minC* | This study |
| LB320 | *minCD::km amyE::spc Pxyl-mGFP-minD(G12V) aprE::cat Pspac-mCherry-minC* | This study |
| LB321 | *minCD::km amyE::spc Pxyl-mGFP-minD(D40A) aprE::cat Pspac-mCherry-minC* | This study |
| LB405 | *minCD::km amyE::spc Pxyl-mGFP-minD* | This study |
| LB406 | *minCD::km amyE::spc Pxyl-mGFP-minD(K16A)* | This study |
| LB407 | *minCD::km amyE::spc Pxyl-mGFP-minD(G12V)* | This study |
| LB408 | *minCD::km amyE::spc Pxyl-mGFP-minD(D40A)* | This study |
| LB409 | *minCD::km minJ::tet amyE::spc Pxyl-mGFP-minD* | This study |
| LB410 | *minCD::km minJ::tet amyE::spc Pxyl-mGFP-minD(K16A)* | This study |
| LB411 | *minCD::km minJ::tet amyE::spc Pxyl-mGFP-minD(G12V)* | This study |
| LB412 | *minCD::km minJ::tet amyE::spc Pxyl-mGFP-minD(D40A)* | This study |
| LB507 | *minD::erm amyE::spc Pxyl-mGFP-minD-BsMTS-tandem(2xAH with linker in between)* | This study |
| LB508 | *minD::erm amyE::spc Pxyl-mGFP-minD-HCV-NS4B-AH* | This study |
| LB559 | *amyE::spc Pxyl-mGFP-minD-HCV-NS4B-AH aprE::cat Pspac-mCherry-minC minCD::km* | This study |
| LB584 | *amyE::spc Pxyl-mGFP-minD-BsMTS-tandem(2xAH with linker in between) aprE::cat Pspac-mCherry-minC minCD::km* | This study |
| FBB043 | *amyE::spc Pxyl-GFP-AH(MinD)* | This study |
| FBB053 | *amyE::spc Pxyl-GFP-BsMTS-tandem(MinD)* | This study |
| FBB046 | *amyE::spc Pxyl-GFP-AH(NS4B)* | This study |
| LB609 | *amyE::spc Phyperspank-sfGFP (from pDR111-N015-sfGFP)* | This study |
| LB643 | *amyE::spc Pxyl-mGFP-minD(I260E) aprE::cat Pspac-mCherry-minC minCD::km* | This study |
| LB644 | *amyE::spc Pxyl-mGFP-minD(D40A I260E) aprE::cat Pspac-mCherry-minC minCD::km* | This study |

**Table S3. Plasmids used in this study**

Antibiotic resistance cassettes are abbreviated as follows: *bla* (ampicillin), *cat* (chloramphenicol), *spc* (spectinomycin).

| Plasmid | Relevant features or genotype | Reference |
|---|---|---|
| pAPNC213 | *bla aprE3' spc Pspac-lacI aprE5'* | (5) |
| pDR111-N015-sfGFP | *bla amyE3' spc Phyperspank-lacI-sfGFP amyE5'* | (6) |
| pHJS113 | *bla amyE3' spc Pxyl-GFP-minD(K16A) amyE5'* | (7) |
| pSG1729 | *bla amyE3' spc Pxyl-GFP amyE5'* | (8) |
| pSG1730 | *bla amyE3' spc Pxyl-GFP-minD amyE5'* | (2) |
| pSG2 | *bla cat* | (9) |
| pSS153 | *bla aprE3' Phag-mCherry-cat aprE5'* | (10) |
| pAPNC*cat* | *bla aprE3' cat Pspac-lacI aprE5'* | This study |
| pHJS112 | *bla aprE3' cat Pspac-mCherry aprE5'* | This study |
| pHJS115 | *bla amyE3' spc Pxyl-GFP-minD(G12V) amyE5'* | This study |
| pHJS116 | *bla amyE3' spc Pxyl-GFP-minD(D40A) amyE5'* | This study |
| pHJS117 | *bla amyE3' spc Pxyl-GFP-MTS(minD) amyE5'* | This study |
| pHJS119 | *bla amyE3' spc Pxyl-GFP-MTS(minD)-tandem(with linker in between) amyE5'* | This study |
| pHJS121 | *bla amyE3' spc Pxyl-GFP-NS4B-AH amyE5'* | This study |
| pHJS123 | *bla amyE3' spc Pxyl-GFP-minD-BsMTS-tandem(with linker in between) amyE5'* | This study |
| pHJS125 | *bla amyE3' spc Pxyl-GFP-minD-NS4B-AH amyE5'* | This study |
| pLB11 | *bla aprE3' cat Pspac-mCherry-minC aprE5'* | This study |
| pLB21 | *bla amyE3' spc Pxyl-mGFP-minD amyE5'* | This study |
| pLB22 | *bla amyE3' spc Pxyl-mGFP-minD(K16A) amyE5'* | This study |
| pLB23 | *bla amyE3' spc Pxyl-mGFP-minD(G12V) amyE5'* | This study |
| pLB24 | *bla amyE3' spc Pxyl-mGFP-minD(D40A) amyE5'* | This study |
| pLB49 | *bla amyE3' spc Pxyl-mGFP-minD-BsMTS-tandem(with linker in between) amyE5'* | This study |
| pLB50 | *bla amyE3' spc Pxyl-mGFP-minD-NS4B-AH amyE5'* | This study |
| pLB71 | *bla amyE3' spc Pxyl-mGFP-minD(I260E) amyE5'* | This study |
| pLB72 | *bla amyE3' spc Pxyl-mGFP(D40A I260E)-minD amyE5'* | This study |

**Table S4. Oligonucleotides used in this study**

| Name | Sequence (5'-3') | Used for / restriction site |
|---|---|---|
| LB1 | GCGCGGGATCCATGAAGACCAAAAAGCAGCAATATG | *minC* / *Bam*HI |
| LB2 | CGCGCGAATTCTCACATTCCTCCCTCAAGCCTTG | *minC* / *Eco*RI |
| HS05 | GCTAATTTTATTGCAATAACAGGTG | To linearize pAPNC213 |
| HS06 | GACCGTTAGCGTTTAAGTACATC | To linearize pAPNC213 |
| HS07 | TAAACGCTAACGGTCCAGTAATATTGACTTTTAAAAAAGG | *cat* cassette |
| HS08 | TGCAATAAAATTAGCTTATAAAAGCCAGTCATTAGGCC | *cat* cassette |
| HS437 | GCGCGGTCGACACATAAGGAGGAACTACTATGGTC | mCherry / *Sal*I |
| HS438 | CGCGCGGATCCTGAGCCGCTTCCTGATTTGTATAATTCGTCCATTCCACC | mCherry / *Bam*HI |
| HS508 | CACAGAATAGTCTTTTAAGTAAGTC | *aprE* |
| HS509 | CCGGAACATCAGGATGCTGAC | *aprE* |
| HS410 | CCTGTCCACACAATCTAAACTTTCGAAAGATCCC | A206K mutation in GFP |
| HS411 | GGGATCTTTCGAAAGTTTAGATTGTGTGGACAGG | A206K mutation in GFP |
| HS112 | GCGGAGTAGGTGCGACAACAACATCTG | K16A mutation in MinD |
| HS113 | CAGATGTTGTTGTCGCACCTACTCCGC | K16A mutation in MinD |
| HS205 | GGAATGATGGCTAAGGAAAAGTCATTTTTCGG | I260E mutation in MinD |
| HS206 | CCGAAAAATGACTTTTCCTTAGCCATCATTCC | I260E mutation in MinD |
| HS320 | CTTCGGGAAAAGTGGGAGTAGGTAAG | G12V mutation in MinD |
| HS321 | CTTACCTACTCCCACTTTTCCCGAAG | G12V mutation in MinD |
| HS322 | GCTTAGTAGATACTGCGATAGGACTGCGC | D40A mutation in MinD |
| HS323 | GCGCAGTCCTATCGCAGTATCTACTAAGC | D40A mutation in MinD |
| FB135 | CTTCGGATCCACGGGCCCCCCCTCGAGTTGGGTGAGGCTATCGTAATAAC | *minD* / *Bam*HI |
| FB136 | CTAGAACTAGAATTCTTAAGATCTTACTCCGAAAAATG | *minD* MTS / *Eco*RI |
| FB149 | CTTCGGATCCACGGATCAGGAGTGCTTGAAGAGCAAAACAAAGG | *minD* MTS / *Bam*HI |
| FB138 | CTAGAACTAGAATTCTTATGATCTCACTCCAAAAAATGATTTAATTTTCGCCATCATTCCTTTGTTTTGTTCTTCCAGCACTCCTCCAGATCTTACTCCGAAAAATGAC | *minD* MTS with overhang generates 2xAH with linker / *Eco*RI |
| FB141 | CTAGAACTAGAATTCTTAAATCCATTGGTGCAGTCTTCTCAGCAGTTGTGTCACTGTCAGTGATGACAGAATGCCGTTTTGCTCTTCAAGCACCTG | HCV-NS4B-AH / *Eco*RI |

**Fig. S1**

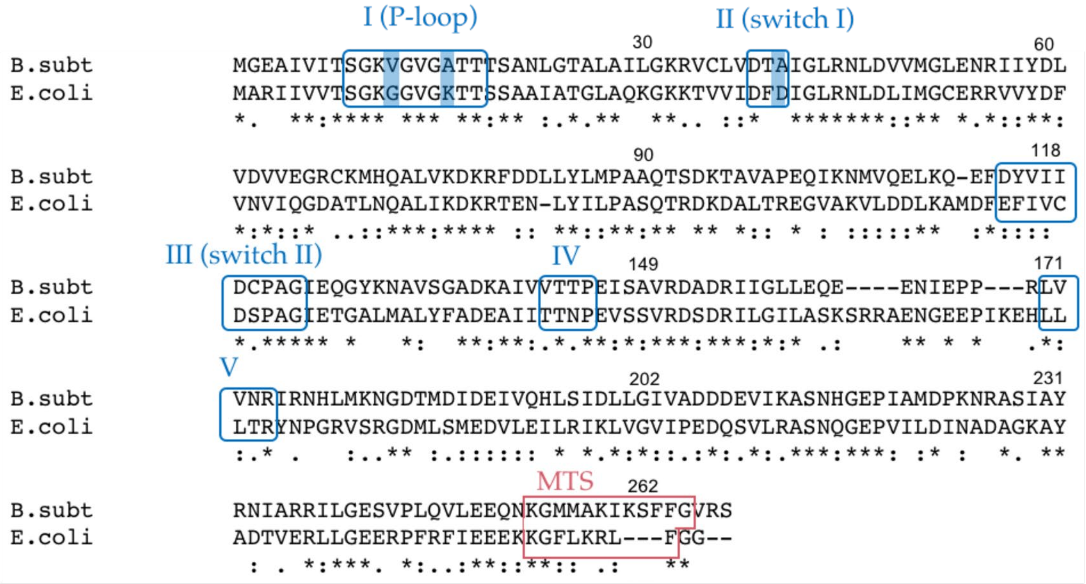


**Fig. S1. Amino acid sequence alignment between *E. coli* and *B. subtilis* MinD proteins**

The *E. coli* and *B. subtilis* MinD proteins share 44 % identical residues. The five conserved motifs of the ATPase protein family are framed in blue (11). The exchanged conserved amino acids are highlighted in blue (G12V, K16A, D40A). The C-terminal membrane targeting amphipathic helices are framed in red (MTS).

**Fig. S2**

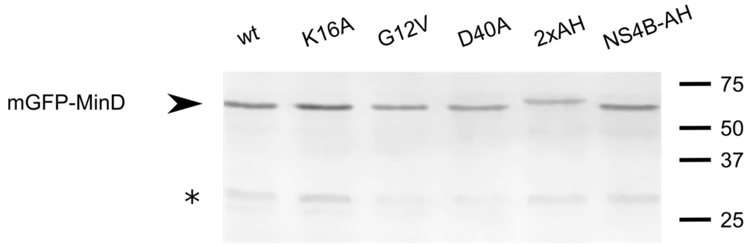


**Fig. S2. Western blot analysis of GFP-MinD variants**

Western blot analysis of strains expressing different mGFP-MinD protein fusions. Cells were grown in LB medium at 37 °C until early exponential phase, followed by induction for 3 doubling times with 0.1 % xylose. Anti-GFP primary antibody was used to detect the different fusion proteins. Strains used: LB305 (wt), LB306 (K16A), LB307 (G12V), LB308 (D40A), LB507 (2xAH), LB508 (NS4B-AH). Black arrowhead indicates full-length mGFP-MinD (~57.7 kDa), the asterisk indicates a nonspecific band.

**Fig. S3**

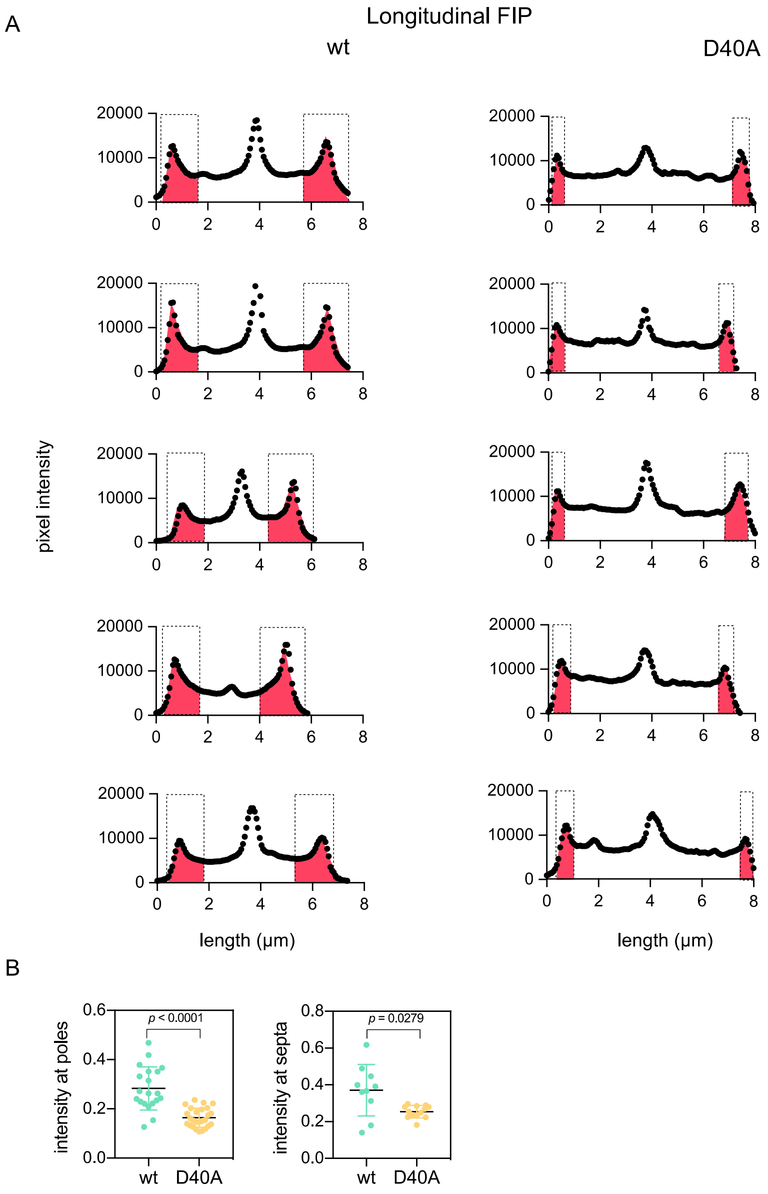


**Fig. S3. Longitudinal fluorescence intensity profiles of GFP-MinD wild type and D40A variants**

(A) Additional examples of longitudinal fluorescence intensity profiles (FIP) of GFP-MinD and GFP-MinD-D40A shown in Fig. 1. (B) Measured pixel intensity, with median values, at poles and septa in longitudinal FIP relative to the intensity of the total integrated area. Significance of difference was confirmed using t-test. The polar and septal areas were selected manually based on the fluorescence gradient. The polar areas are indicated in red.

**Fig. S4.**

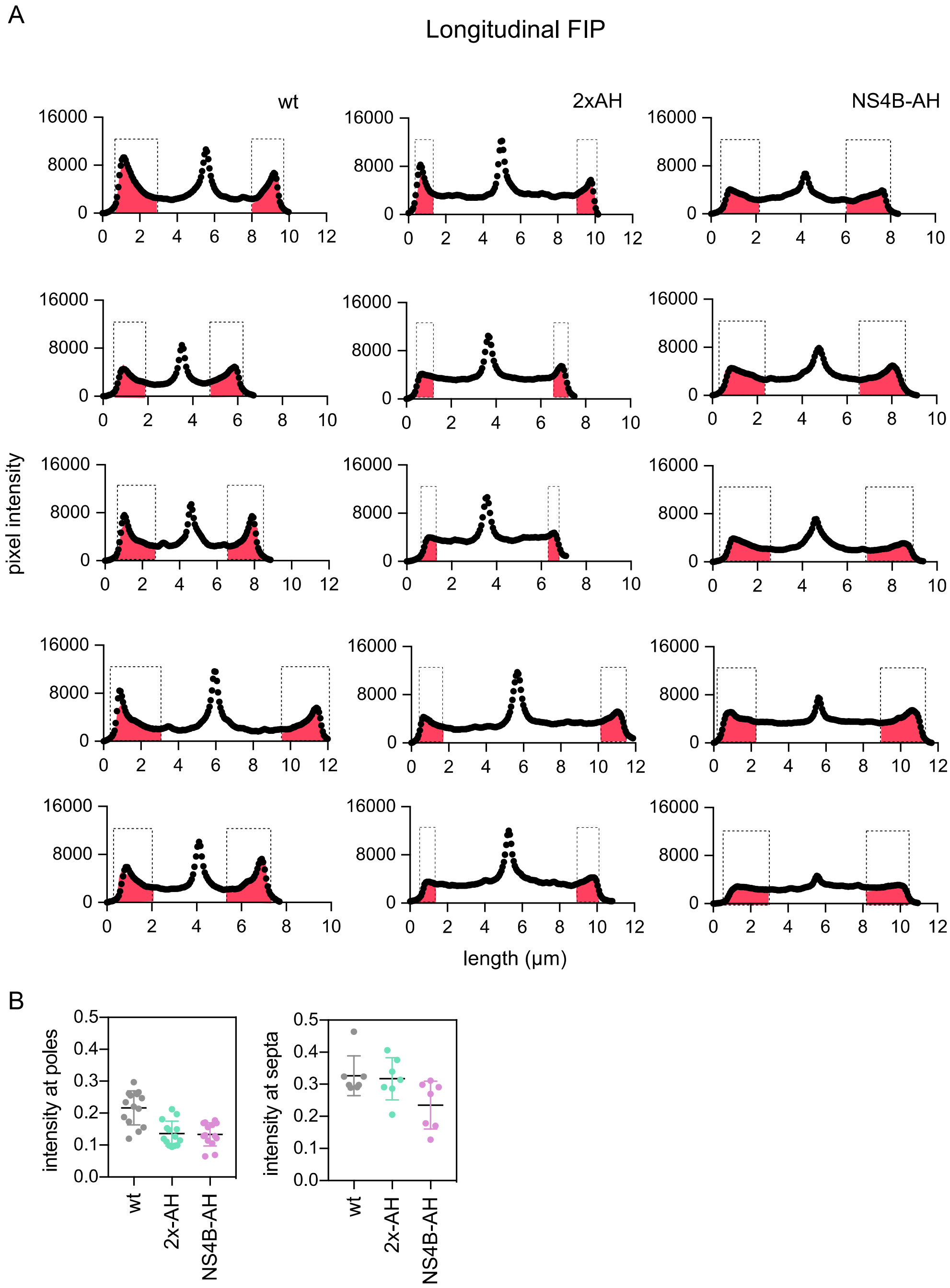


**Fig. S4. Longitudinal fluorescence intensity profiles of mGFP-MinD with different membrane anchors.**

(A) Additional examples of longitudinal fluorescence intensity profiles (FIP) from cells expressing mGFP–MinD variants with different membrane affinities shown in Fig. 5. (B) Pixel intensity, and median values, at poles and septa in longitudinal FIP relative to the integrated area of intensity. The polar and septal areas were selected manually based on the fluorescence gradient. The polar areas are indicated in red.

**Fig. S5.**

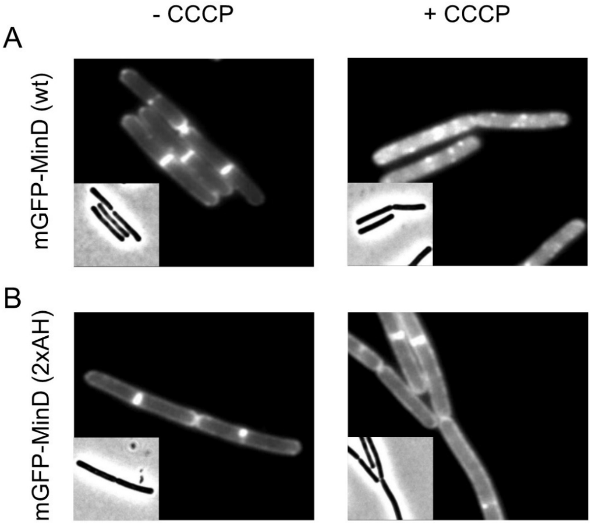


**Fig. S5. Effect of membrane potential dissipation on membrane binding**

Fluorescence images of cells expressing mGFP–MinD variants with the wild type amphipathic helix (wt) (A), and the strong (2xAH) membrane binding amphipathic helix (B). Corresponding phase contrast images are shown in the insets. The membrane potential was dissipated by the proton-ionophore CCCP (100 μM for 10 min). mGFP-MinD fusions were expressed from the ectopic *amyE* locus using a xylose-inducible promoter and a Δ*minD* background. Strains used: (A) LB249, (B) LB507.

**Fig. S6.**

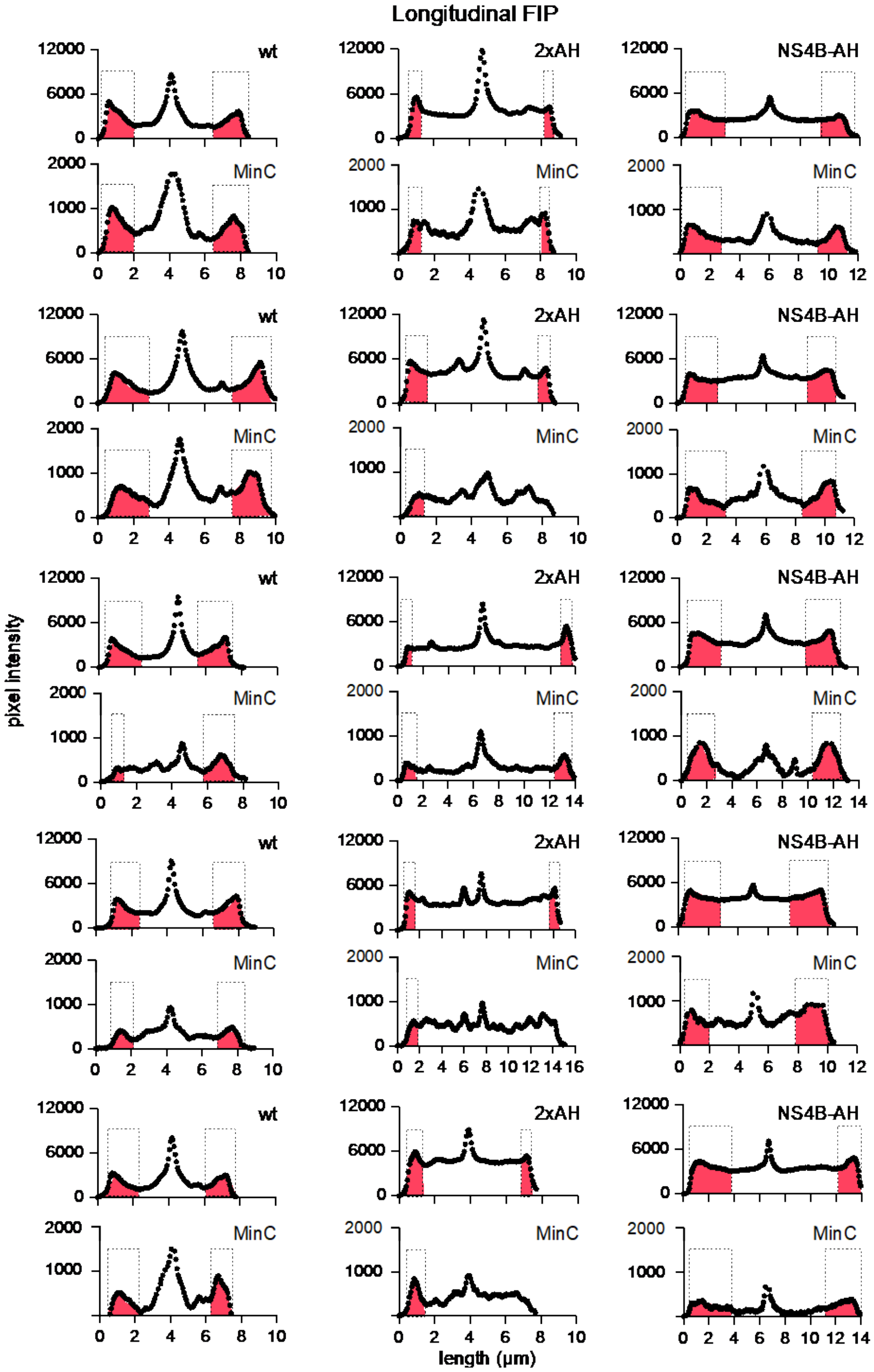


**Fig S6. Longitudinal fluorescence intensity profiles of mCherry-MinC in cells expressing different mGFP-MinD membrane-binding variants**

Additional examples of longitudinal fluorescence intensity profiles (FIP) from cells expressing mCherry-MinC and mGFP–MinD variants with different membrane affinities shown in Fig. 7. Upper FIPs and lower FIPs depict MinD and MinC gradients, respectively. The latter is indicated by “MinC”.
